# Effects of long-term fasting: longitudinal epigenetic responses in humans

**DOI:** 10.64898/2026.06.12.731903

**Authors:** Siyuan Luo, Marie Knufinke, Cassandre Kinnaer, Laurence Chapatte, Sébastien Nusslé, Semira Gonseth Nusslé, Franziska Grundler, Francoise Wilhelmi de Toledo, Steve Horvath, Robin Mesnage, Ferdinand von Meyenn

## Abstract

Prolonged fasting induces marked metabolic adaptations, but whether such time-limited yet intensive interventions engage molecular processes related to biological aging in humans remains unclear. To address this, we investigated physiological and epigenetic responses to a 12-day medically supervised fasting intervention with longitudinal follow-up in 32 participants. Fasting elicited coordinated systemic physiological changes across metabolic and hematological parameters, with partial persistence at one month. Genome-wide DNA methylation analyses revealed modest but detectable CpG-level changes, primarily emerging at follow-up and distributed across genomic contexts. In parallel, epigenetic aging clocks showed clock-specific and time-dependent responses, with substantial inter-individual variability. Lower baseline epigenetic age acceleration was consistently associated with greater fasting-induced weight loss, suggesting a link between epigenetic state and metabolic responsiveness.

Together, these findings indicate that prolonged fasting induces coordinated physiological adaptations alongside structured changes in epigenetic aging measures that are not captured by conventional clinical biomarkers. This study highlights epigenetic clocks as integrative molecular readouts for probing aging-related responses to short-term metabolic interventions, while also delineating their current interpretive limits.

## Introduction

Fasting and caloric restriction have been shown to extend lifespan and support healthy aging in a wide range of model organisms, including yeast, rodents, and non-human primates^1–3^. In humans and non-human primates, energy-restriction and fasting regimens have been associated with improvements in cardiometabolic health and risk factors for age-related disease, such as glucose control, lipid profiles, and blood pressure^4–7^. Together, these findings motivate the investigation of fasting as a potential intervention for improving lifespan- and healthspan-related outcomes in humans.

Mechanistically, fasting induces coordinated metabolic and cellular adaptations that intersect with core pathways implicated in aging biology. At the systemic level, fasting promotes a metabolic shift from glucose to lipid oxidation, accompanied by increased circulating ketone bodies such as β-hydroxybutyrate^8^. At the cellular level, this shift in nutrient availability activates energy-sensing pathways including AMPK-and SIRT1-dependent signaling, which regulate autophagy, mitochondrial function, oxidative stress responses, and genome maintenance^7,9–12^. These processes play central roles in established models of aging and age-related functional decline^13,14^.

While mechanistic data from animal models are increasingly available^3^, controlled studies examining fasting-induced aging-related adaptations in humans remain comparatively limited. Human fasting studies are often constrained by heterogeneity in fasting duration, caloric intake, adherence, and concurrent lifestyle factors^15^.

Medically supervised clinical fasting programs provide a more standardized experimental context, with defined dietary exposure, structured daily routines, and systematic clinical monitoring^5,16^. Within such settings, prolonged inpatient fasting protocols enable repeated biological sampling under controlled conditions, facilitating the investigation of both acute responses during fasting and post-intervention dynamics following food reintroduction.

Despite extensive characterization of fasting-induced changes in body weight, cardiometabolic markers, and inflammatory profiles^5,15^, it remains unclear whether time-limited but intensive fasting interventions engage biological processes associated with longer-term aging trajectories. Conventional physiological and biochemical biomarkers primarily reflect acute metabolic or inflammatory states and may not capture slower, integrative remodeling processes that unfold over extended timescales. This raises the question of whether alternative molecular readouts are required to assess aging-related responses to fasting in humans.

DNA methylation–based epigenetic clocks quantify age-associated methylation patterns and are widely used as biomarkers of biological aging^17,18^. These measures show robust associations with long-term morbidity and mortality risk across multiple cohorts^19^. In contrast to many conventional physiological biomarkers, epigenetic age measures are thought to reflect cumulative molecular and cellular processes, including changes in cell turnover, lineage composition, and regulatory state^20,21^. As a result, epigenetic aging may capture aspects of biological aging that are less sensitive to transient physiological perturbations and more reflective of sustained or integrative adaptation.

Epigenetic clocks differ substantially in their training objectives and modeling strategies^22^. First-generation clocks were trained primarily to predict chronological age and are not optimized for health stratification^23–25^. In contrast, later-generation clocks aim to capture inter-individual differences in biological aging, mortality risk, or longitudinal deterioration of multisystem health. Some clocks derive latent aging constructs from clinical or longitudinal biomarkers and subsequently regress DNA methylation patterns onto these constructs, as exemplified by PhenoAge^26^, SystemsAge^27^, and DunedinPACE^28^. Others model DNA-methylation–based proxies of health-related exposures or biomarkers and integrate these proxies to predict time-to-death, as in GrimAge^29^ and GrimAge v2^30^. Several clocks expose their constituent components, enabling modular analyses and improved interpretability^27,29,30^. More recently, causality-informed approaches such as CausalAge and its components DamAge and AdaptAge aim to disentangle damaging and adaptive aging processes by using genetic variation to inform causal relationships between DNA methylation and aging-related traits^31^.

Alongside conceptual advances, methodological refinements have improved the robustness of epigenetic clocks in longitudinal settings. Principal-component–based implementations were developed to enhance test–retest reliability by aggregating correlated CpGs into stabilized components while preserving original training targets^32^. More recent clocks, including GrimAge v2^30^ and DunedinPACE^28^, were explicitly designed to incorporate CpGs with high technical reliability, improving reproducibility by construction. These developments are particularly relevant for short-term intervention studies, where biological effects may be modest relative to technical variability.

Most epigenetic clocks have been validated primarily for their ability to predict current or future morbidity and mortality. In contrast, their responsiveness to interventions targeting aging-related biology remains less well characterized. Existing dietary intervention studies have reported heterogeneous epigenetic aging responses, with different interventions affecting distinct subsets of clocks^33–37^. Notably, closely related interventions have produced discordant results across clocks, and responsiveness does not appear to be uniform^38^. To date, epigenetic aging dynamics have not been directly examined in the context of prolonged clinical fasting.

In this study, we investigate the effects of a 12-day medically supervised fasting intervention followed by a structured food reintroduction phase on multiple epigenetic clocks capturing distinct aspects of aging biology. Whole-blood samples were collected at baseline (day -1), during fasting (days +4 and +10), and one month after fasting completion. The one-month follow-up was chosen to assess epigenetic aging dynamics after resolution of the acute fasting response and return to metabolic steady state under habitual dietary conditions. In parallel, an extensive panel of clinical and biochemical measurements was obtained throughout the intervention period. This design enables evaluation of whether fasting induces detectable changes in epigenetic measures of biological aging during and after the intervention, and whether such changes capture fasting-induced adaptations beyond those reflected by routine clinical markers, thereby informing the utility of epigenetic clocks as candidate surrogate endpoints in short-term aging intervention studies.

## Methods

### Study design

This was a prospective, monocentric, single-arm interventional study conducted at the Buchinger Wilhelmi Clinic (Überlingen, Germany). 32 healthy adults (16 women and 16 men) participated in a medically supervised prolonged fasting intervention lasting 12 days.

The intervention followed a standardized Buchinger fasting protocol^5^. Participants began with a transition day (D0) involving a calorie-restricted vegetarian diet (∼600 kcal) to ease the metabolic shift into fasting. This was followed by 12 consecutive days of fasting (D+1 to D+12), during which daily caloric intake was approximately 200–250 kcal, provided as organic fruit juices, vegetable broths, and honey, alongside ≥2.5 L/day of fluids (water and herbal teas). The protocol also included bowel cleansing procedures (initial sodium sulfate administration and periodic enemas), structured rest periods, and up to 3 hours per day of light physical activity (e.g., walking, stretching).

Following the fasting period, participants underwent a standardized four-day food reintroduction phase (D+13 to D+16) with an ovo-lacto-vegetarian diet and a gradually increasing caloric intake. After discharge, participants resumed an unrestricted, self-selected diet. A clinical follow-up visit was conducted one month after the end of fasting to assess medium-term effects.

### Sample collection

Baseline assessments were performed upon arrival at the clinic prior to initiation of fasting (D−1), under habitual dietary conditions. Venous blood samples were collected in the morning at four time points: baseline (D-1), day 4 of fasting (D+4), day 10 of fasting (D+10), and one month post-fasting (M+1). DNA methylation analyses were performed on samples collected at D−1, D+10, and M+1.

During the inpatient stay, daily measurements included anthropometric measurements (body weight, waist circumference and height), vital signs (blood pressure and heart rate measured using the boso medicus exclusive (BOSCH + SOHN GmbH u. Co. KG, Jungingen, Germany), capillary blood samples for glucose and β-hydroxybutyrate using the GlucoMen areo 2K (Berlin-Chemie AG, Berlin, Germany), as well as self-reported wellbeing and symptom scores. At the one-month follow-up (M+1), anthropometric measurements and self-reported scores were repeated.

Baseline demographic characteristics, medical history, and lifestyle factors were assessed at D-1 (Table 1). Medical diagnoses were derived from baseline clinical assessments and medical history and harmonized across overlapping ICD-10 codes into clinically meaningful categories for descriptive analyses. Overweight or obesity was defined as BMI ≥ 25 kg/m^2^. Elevated blood pressure or hypertension was defined based on documented borderline or established arterial hypertension.

**Table 1:** Baseline demographic, lifestyle, and clinical characteristics of the GENESIS study cohort. Values are presented as mean ± SD (range) or number (percentage), as appropriate. Medical diagnoses were harmonized across overlapping ICD-10 codes into clinically meaningful categories. Detailed definitions and assessment procedures are provided in the Methods section.

| Characteristic | Total cohort (n = 32) |
| --- | --- |
| **Demographics** |  |
| Age, years | 47.88 ± 12.14 (25 - 72) |
| Sex, n (%) |  |
| Male | 16 (50%) |
| Female | 16 (50%) |
| Ethnicity, n (%) |  |
| European ancestry | 31 (96.9%) |
| South Asian ancestry | 1 (3.1%) |
| **Medical Diagnoses** |  |
| Overweight or obesity (BMI ≥ 25 kg/m<sup>2</sup>), n (%) | 14 (44%) |
| Elevated blood pressure / hypertension, n (%) | 2 (6%) |
| Impaired glucose metabolism, n (%) | 2 (6%) |
| Hypercholesterolaemia, n (%) | 1 (3%) |
| **Anthropometrics** |  |
| Body weight, kg | 79.16 ± 13.01 |
| Body mass index (BMI), kg/m<sup>2</sup> | 26.14 ± 3.92 (21.00 - 37.40) |
| **Vital signs** |  |
| Systolic blood pressure, mmHg | 124.19 ± 15.40 |
| Diastolic blood pressure, mmHg | 79.65 ± 10.69 |
| Resting heart rate, bpm | 67.09 ± 13.80 |
| **Lifestyle factors** |  |
| Smoking status, n (%) |  |
| No smoking | 14 (43.75%) |
| Former smoking | 18 (56.25%) |
| Alcohol consumption, standard drinks/week | 6.05 ± 5.39 |
| Leisure-time physical activity score (0–100) | 36.70 ± 26.55 |

Impaired glucose metabolism was defined by the presence of prediabetes and/or insulin resistance. Hypercholesterolaemia was recorded based on prior clinical diagnosis. Lifestyle factors, including smoking status, alcohol consumption, and leisure-time physical activity, were assessed by self-report using standardized questionnaires at baseline. Smoking status was categorized as no smoking or former smoking. Current smoking as well as regular medication use were exclusion criteria. Alcohol intake was expressed as average standard drinks per week. Leisure-time physical activity was summarized using a composite activity score ranging from 0 to 100, with higher values indicating greater activity^39^.

### Sample processing and data generation

Venous whole blood was collected in a fasted state in the morning at four time points (D–1, D+4, D+10, and M+1) into EDTA-coated tubes and stored at –80 °C. Genomic DNA was extracted using the Quick-DNA™ 96 Plus Kit (Zymo Research, D4071) following the manufacturer’s protocol. DNA concentration and purity was assessed by absorbance using the Varioskan (Thermo Scientific) fitted with a µDrop™ Duo Plate (Thermo Scientific, N12391M2), and DNA was normalized to 16ng.µl-1. The samples underwent an overnight bisulfite conversion and were purified using the EZ-96 DNA Methylation™ Kit (Zymo Research, D5004) kit. Samples proceeded directly to the Illumina Infinium HD assay (Illumina Methylation Epic Array Cat. No.

WG-317-1001) preparation. Briefly, the samples go through overnight whole genome amplification at 37°C. The samples are then fragmented for 1 hour at 37°C, precipitated at 4°C for 30 minutes, and pelleted at 4°C for 20 minutes at 3000g. After drying at room temperature, the samples are re-suspended in RA1 at 48°C for 1 hour. Samples are then denatured at 95°C for 20 minutes. After a brief cool down, they are hybridized overnight at 48°C in Illumina Methylation EPIC Array. The following day arrays are washed, stained and sealed then read on Illumina’s iScan array scanner. Raw intensity data files were processed using the preprocessing pipeline described below.

### DNA methylation analysis

#### Preprocessing

DNA methylation data generated using the Illumina Infinium MethylationEPIC array were imported, processed, and quality-controlled using the R/Bioconductor packages Meffil v1.3.6^40^ and SeSAMe v1.18.4^41^. Sample-level quality control (QC) was performed using Meffil, including inference of biological sex from methylation patterns and comparison with reported sex; no discrepancies were observed. Additional sample-level QC metrics included assessment of control probes, detection p-values, and bead counts, following Meffil’s recommended procedures. Probe-level QC was conducted using a combination of Meffil and SeSAMe. Probes were first filtered using Meffil to remove those with detection p-values > 0.01 in more than 10% of samples or with bead counts < 3 in more than 10% of samples. Subsequently, SeSAMe was used to exclude probes failing built-in quality masks and to mask probes identified as unreliable based on the out-of-band signal distribution using the pOOBAH() function (p-value threshold = 0.05). Following probe filtering, channel inference, background subtraction, and non-linear dye-bias correction were performed using SeSAMe according to the package documentation.

#### Batch correction

DNA methylation array experiments were conducted across multiple dates. To correct for potential batch effects in methylation beta values, we applied the ComBat method^42^ using the champ.runCombat() function from R/Bioconductor package ChAMP v2.38.0^43^, specifying batchname=c(“Date_Scan”). Batch-corrected beta values were used for downstream epigenetic age prediction analyses.

For differential methylation analyses, batch correction was not applied. Instead, uncorrected beta values were used, with scan date (Date_Scan) included as a covariate in the linear models to account for potential confounding effects (see below for details).

#### Differential methylation analysis

Differentially methylated CpG sites were identified using the R/Bioconductor package limma v3.56.2^44^. Beta values were first converted to M-values to improve the statistical properties of the data. Differential methylation was assessed using linear models with a correlation structure for repeated measures, implemented via the duplicateCorrelation framework in limma. For each CpG (M-value), methylation level was modeled as the outcome, with time point as the main predictor (fixed effect).

Additional covariates included scan date (Date_Scan), chronological age, sex, and measured lymphocyte proportions.

Because whole-blood DNA methylation profiles are sensitive to white blood cell composition, lymphocyte proportion was included to account for blood cell mixture–related variability. Principal component analysis of hematological parameters identified lymphocyte and neutrophil proportions as the dominant source of between-sample variance (Supplementary Figure 4). However, these variables were strongly inversely correlated (Pearson r=-0.93), and therefore only lymphocyte proportion was included to avoid multicollinearity. Mean levels of these cell-type proportions did not show statistically significant changes across timepoints, reducing the likelihood of over-adjustment of fasting-associated time effects.

Inter-individual correlation arising from repeated measurements was accounted for by specifying participant ID as the grouping variable. Contrasts were constructed to compare the D+10 and M+1 time points against the baseline (D−1). CpG sites with a false discovery rate (FDR) < 0.05 and an absolute change in beta values > 0.02 were considered significantly differentially methylated.

KEGG pathway enrichment analysis was performed using the gometh() function from the R/Bioconductor package missMethyl v1.42.0^45^.

#### Epigenetic clock calculation

Epigenetic age and pace-of-aging measures were computed using multiple established DNA methylation–based clocks across the three time points.

DunedinPACE predictions were generated using the PACEProjector() function from the R package DunedinPACE (v0.99.0), obtained from the GitHub repository danbelsky/DunedinPACE. Predictions for first- and second-generation epigenetic clocks in their principal component (PC) implementations—including Horvath 1 (pan-tissue), Horvath 2 (skin and blood), Hannum, PhenoAge, GrimAge, and DNAm-derived telomere length (DNAmTL)—were computed using the R scripts provided in the original publication of PC clocks^32^, available from the GitHub repository MorganLevineLab/PC-Clocks. Original GrimAge and GrimAge2 predictions were computed using in-house scripts. Causality-enriched and system-level clocks, including AdaptAge, DamAge, CausAge, and SystemsAge, were computed using the R package methylCIPHER (v0.2.0), obtained from the GitHub repository HigginsChenLab/methylCIPHER.

### Statistical analysis

#### Analysis of clinical parameters

Principal Component Analysis (PCA) was used to characterize multivariate structure in clinical and hematological parameters. PCA was performed on a predefined set of core blood and clinical parameters (Table 2), after excluding variables with missing values. Three PCAs were conducted:(1) PCA of all blood parameters at baseline (D-1; Figure 2b & Supplementary Figure 2); (2) PCA of change-from-baseline values at D+10 and M+1 (Figure 2c & Supplementary Figure 3); and (3) PCA of white blood cell composition parameters across all three time points (D-1, D+10, and M+1; Supplementary Figure 4). Variables were centered and scaled prior to analysis. PCA was conducted using the R/Bioconductor package PCAtools v2.20.0.

**Table 2:**
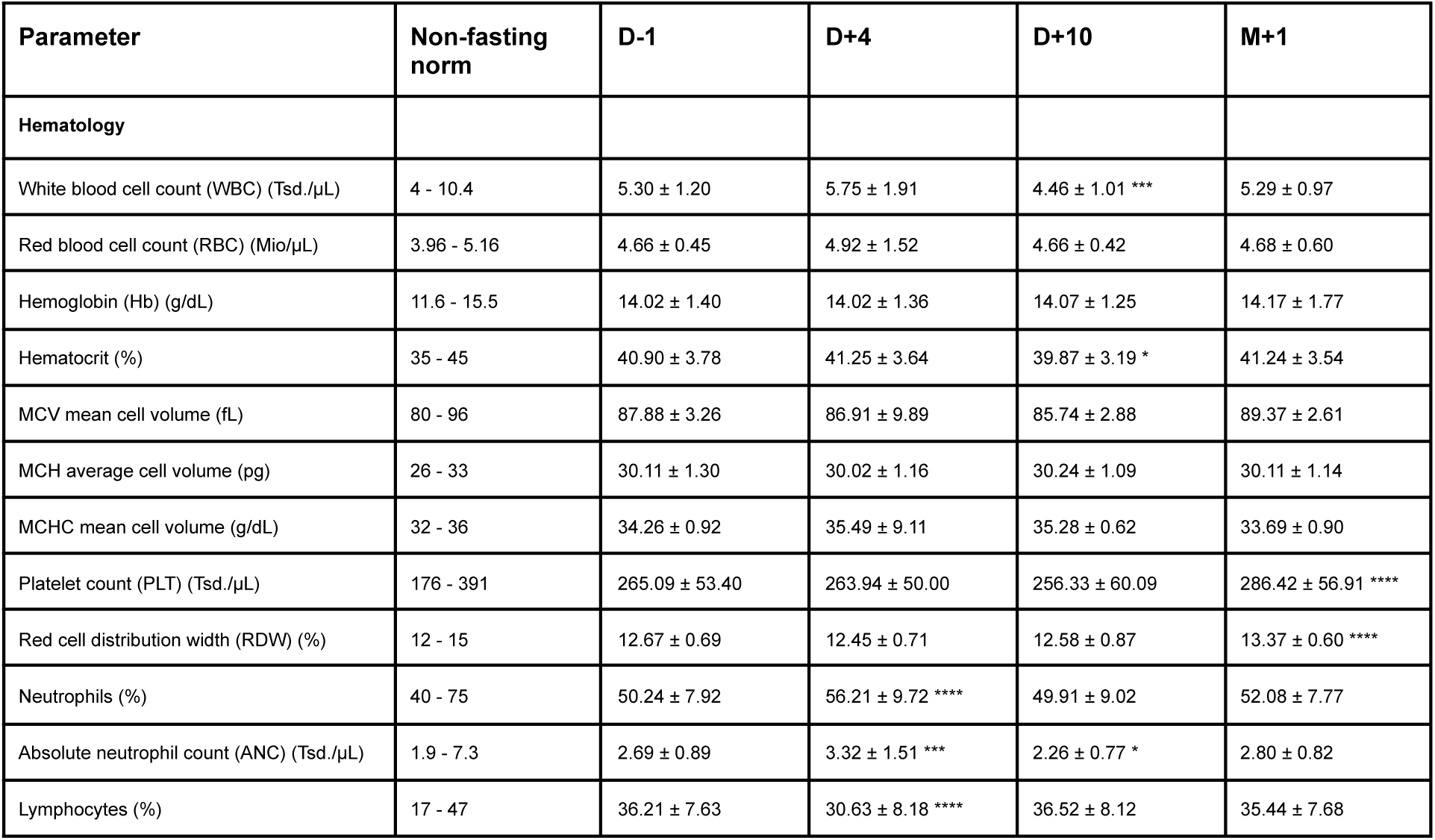
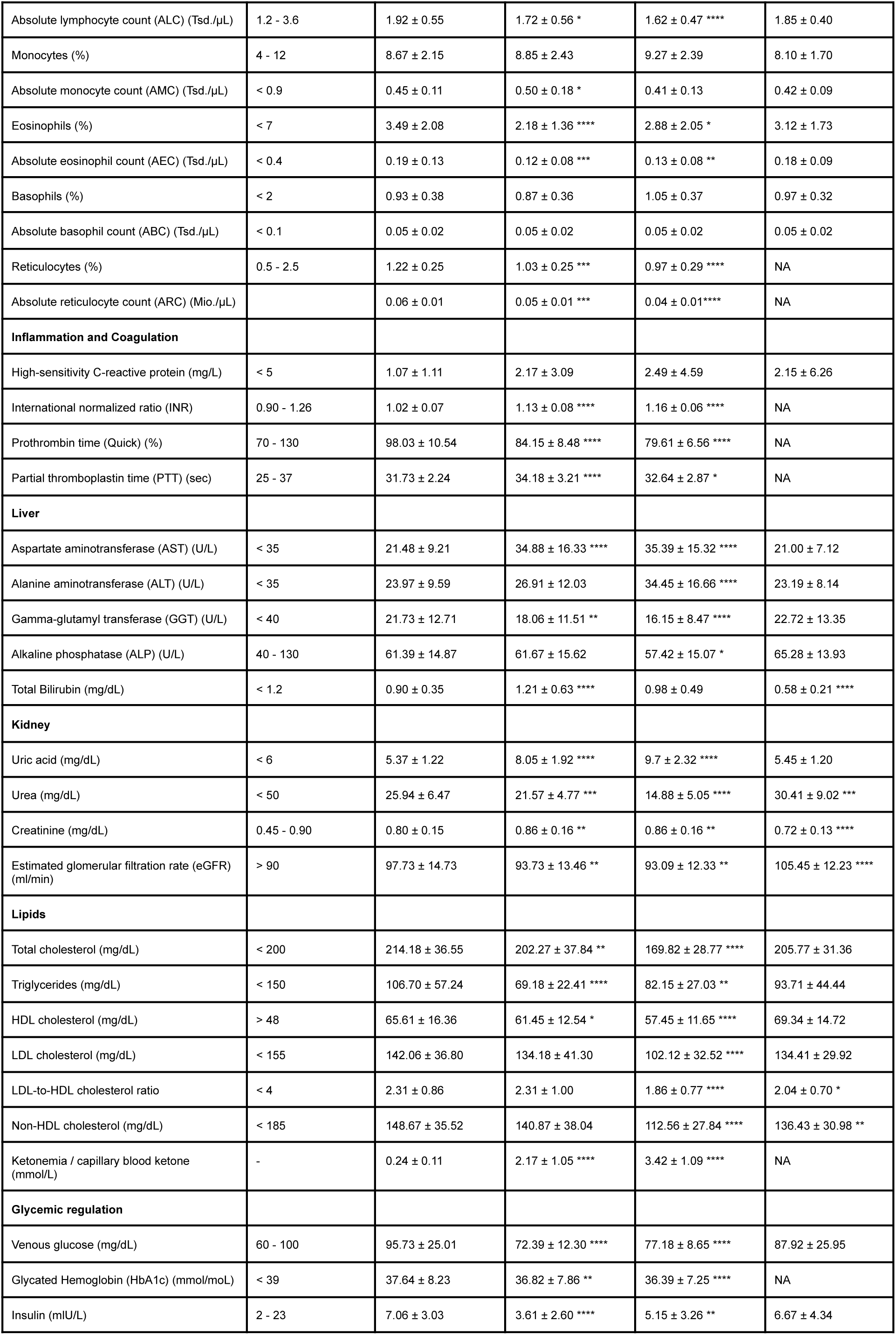
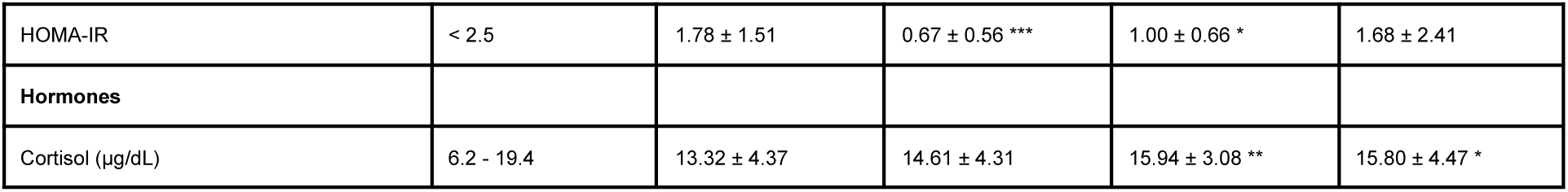
Blood parameters measured at D-1, D+4, D+10 and M+1. Values are presented as mean ± SD. Significance levels for comparisons with D–1 are indicated as follows: * = p<0.05, ** = p<0.01, *** = p<0.001. **** = p<0.0001. Not all parameters were measured at M+1 (indicated with NA).

Longitudinal changes in individual clinical parameters (Figure 2d; Table 2, Supplementary Table) were analysed using linear mixed-effects models (LMMs) to account for repeated measurements within participants. Time point was modelled as a categorical fixed effect, and participant ID was included as a random intercept to capture subject-specific baseline differences. Models were fitted using restricted maximum likelihood, and pairwise contrasts between time points were used to assess phase-specific differences without assuming linear temporal trends. P-values were adjusted for multiple testing where indicated. Analyses were performed using the R package lme4 (v1.1). Full results are provided in Table 2 and Supplementary Table.

#### Analysis of epigenetic clocks

##### Residual age acceleration and longitudinal modeling

For each epigenetic clock, residual age acceleration (RAA) was calculated at three time points by regressing epigenetic age (or pace/telomere-derived measures) on chronological age and extracting the residuals. RAA values were standardized by the baseline (D-1) standard deviation of each clock. Change-from-baseline standardized residual age acceleration (ΔSRAA) was then computed at D+10 and M+1 for each individual. This standardization places clock responses on a common scale, enabling direct comparison across clocks.

Within-subject changes in individual epigenetic clocks were assessed using linear mixed-effects models. For each clock, time point was included as a fixed effect, along with sex and hematological covariates (lymphocyte proportions). Participant ID was included as a random intercept to account for repeated measurements.

Primary longitudinal analyses are shown in Figure 4, with additional details provided in Supplementary Information Section 1.

##### Sensitivity to blood cell composition adjustment

To evaluate the sensitivity of clock time effects to blood cell composition adjustment, linear mixed-effects models were additionally fitted without lymphocyte adjustment. Standardized time-effect estimates obtained from adjusted and unadjusted models were compared across clocks and time points (Supplementary Figure 7, 10, and 12). Additional interpretation is provided in Supplementary Information Section 1.

##### Decomposition analysis

To further assess the contribution of blood cell composition to fasting-associated clock dynamics, we decomposed each standardized clock signal into a lymphocyte-associated component and a residual component. For each clock, a linear mixed-effects model was fitted with standardized residual age acceleration (SRAA) as the outcome, lymphocyte proportion as a fixed effect, and participant ID as a random intercept:

clock_SRAA ∼ Lymphocyte_proportion + (1 | patient_ID)

The lymphocyte-associated component was defined as the fixed-effect prediction from this model, obtained using predict(model, re.form = NA), thereby excluding participant-specific random intercepts. The residual component was then calculated as the observed standardized clock value minus the lymphocyte-associated component.

To test whether each component changed over time, we fitted separate linear mixed-effects models for the lymphocyte-associated and residual components:

composition_component ∼ Timepoint + Sex + (1 | patient_ID)

residual_component ∼ Timepoint + Sex + (1 | patient_ID)

This analysis was used to distinguish clock changes associated with lymphocyte composition from residual, composition-independent clock dynamics. Decomposition results are presented in Supplementary Figure 9, 11, and 13, with additional interpretation provided in Supplementary Information Section 1.

##### Principal component analysis of clock responses

To characterize shared structure across clock responses, PCA was performed separately at D+10 and M+1 using participant-level vectors of ΔSRAA across clocks (Supplementary Figure 14). PCA was conducted on centered and scaled ΔSRAA values using the prcomp() function in R. As a robustness check, PCA was repeated after excluding high-variance clocks (DamAge) to assess the stability of dominant components.

To interpret the biological relevance of PCA-defined response axes, participant scores for the leading principal components (PC1–PC3) were tested for association with pre-specified panels of clinical parameters capturing metabolic state, stress, hematological measures, and subjective well-being. Variables were selected to represent broad physiological domains rather than exhaustive biomarker panels.

Associations were assessed using Spearman rank correlation, and p-values were adjusted for multiple testing within each panel using the Benjamini–Hochberg procedure. Some results are shown in Supplementary Figure 15 and 16. Full results are provided in Supplementary Table 4.

##### Associations with fasting responsiveness

Associations between epigenetic age acceleration and maximum weight loss percentage were assessed using linear mixed-effects models with maximum weight loss percentage as the primary predictor. Fixed-effect covariates included time point, sex, and lymphocyte proportions, with participant ID included as a random intercept. Marginal effects of maximum weight loss percentage at each time point were estimated using the R package ggeffects v2.2.1.

##### Sensitivity of weight loss associations to baseline BMI

To assess potential confounding by adiposity, sensitivity analyses were performed by additionally including baseline BMI as a fixed-effect covariate. Additional interaction models included a maximum weight loss percentage × baseline BMI interaction term to evaluate whether associations differed across adiposity levels. Complementary baseline-only sensitivity analyses were performed using linear regression models with maximum weight loss percentage as the outcome and baseline residual age acceleration (RAA) as the primary predictor. Sex and baseline BMI were included as covariates to account for potential confounding by sex-related and adiposity-related differences in weight loss responses.

Additional R packages used for statistical analyses include lme4 v1.1.37, lmerTest v3.1.3 and emmeans v1.11.1.

### Visualization

R v4.5.0 and ggplot2 v3.5.2 were used to analyze data and generate plots.

## Results

### Study cohort

The GENESIS study is a prospective, monocentric, single-arm interventional trial conducted at the Buchinger Wilhelmi clinics in Germany^16^. The study design and data collection schedule are summarized in Figure 1, with methodological details provided in the Methods section. The GENESIS cohort has previously been used to investigate fasting-induced adaptations in skeletal muscle^46^, autonomic nervous system function^47^, erythrocyte fatty acid membrane composition^48^, and HDL cholesterol efflux capacity^49^.

**Figure 1:**
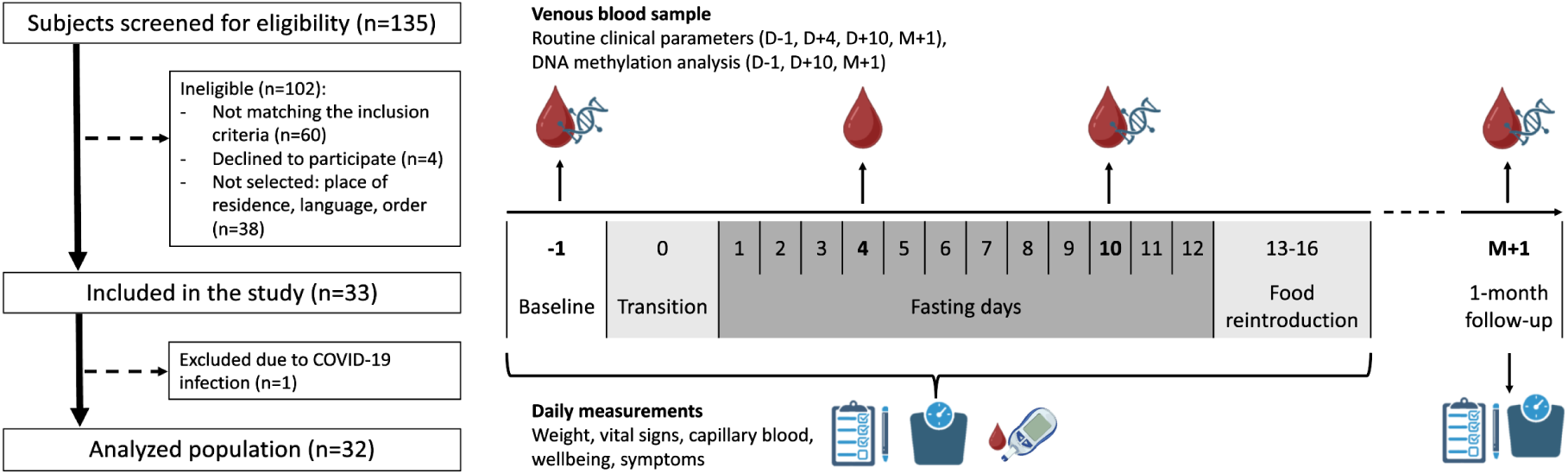
Study flow and measurement schedule for the GENESIS trial. The left panel summarizes participant recruitment and study flow, and the right panel shows the timing of sample collection and data generation. Participants underwent screening and baseline assessment one day before the start of the fasting protocol (D-1), followed by a transition day (D0) and a 12-day medically supervised fasting period (D+1 to D+12), a structured food reintroduction phase (D+13 to D+16), and a one-month follow-up visit (M+1). Routine clinical measurements, including anthropometric parameters, vital signs, capillary blood measurements, and symptom assessments, were collected daily during the inpatient stay and at M+1. Venous blood samples were obtained at D-1, D+4, D+10, and M+1, with DNA methylation profiling performed on samples collected at D-1, D+10, and M+1.

A total of 135 individuals were screened for eligibility, of whom 33 were included in the study. One person was excluded due to an acute SARS-CoV-2 infection, as defined in the termination criteria, leaving 32 participants in the final cohort. All participants contributed baseline clinical data and completed the 12-day medically supervised fasting intervention and participated in the one-month follow-up assessments. Longitudinal clinical measurements were available for all participants, with limited missingness in individual clinical parameters. Venous blood samples for DNA methylation analysis were collected at baseline (D−1), during fasting (D+10), and at the one-month follow-up (M+1). All 96 DNA methylation profiles generated using the Illumina EPIC array passed quality control (see Methods) and were included in downstream analyses.

The study cohort comprised 16 male participants (aged 31-63 years) and 16 female participants (aged 25-72 years), with a mean age of 47.9 years. Baseline body mass index (BMI) ranged from 21.0 to 37.4 kg/m^2^ (mean 26.1 kg/m^2^; Supplementary Figure 1). The cohort exhibited heterogeneity in baseline cardiometabolic risk profiles. All participants were non-smokers and regular medication use was an exclusion criterion. Alcohol consumption and leisure-time physical activity varied across individuals as assessed by standardized questionnaires. Baseline demographic, lifestyle, and longitudinal clinical parameters are summarized in Tables 1, 2 and Supplementary Table 1.

### Systemic physiological changes during and after fasting

All 32 participants showed robust engagement with the fasting intervention, reflected by changes in body weight and alterations in metabolic parameters consistent with canonical fasting-associated energy metabolism^5,50^ (Figure 2a,d; Table 2, Supplementary Table 1). Although some parameters transiently deviated from standard non-fasting reference ranges during fasting (Table 2), these changes were generally consistent with expected physiological adaptations to prolonged fasting.

**Figure 2:**
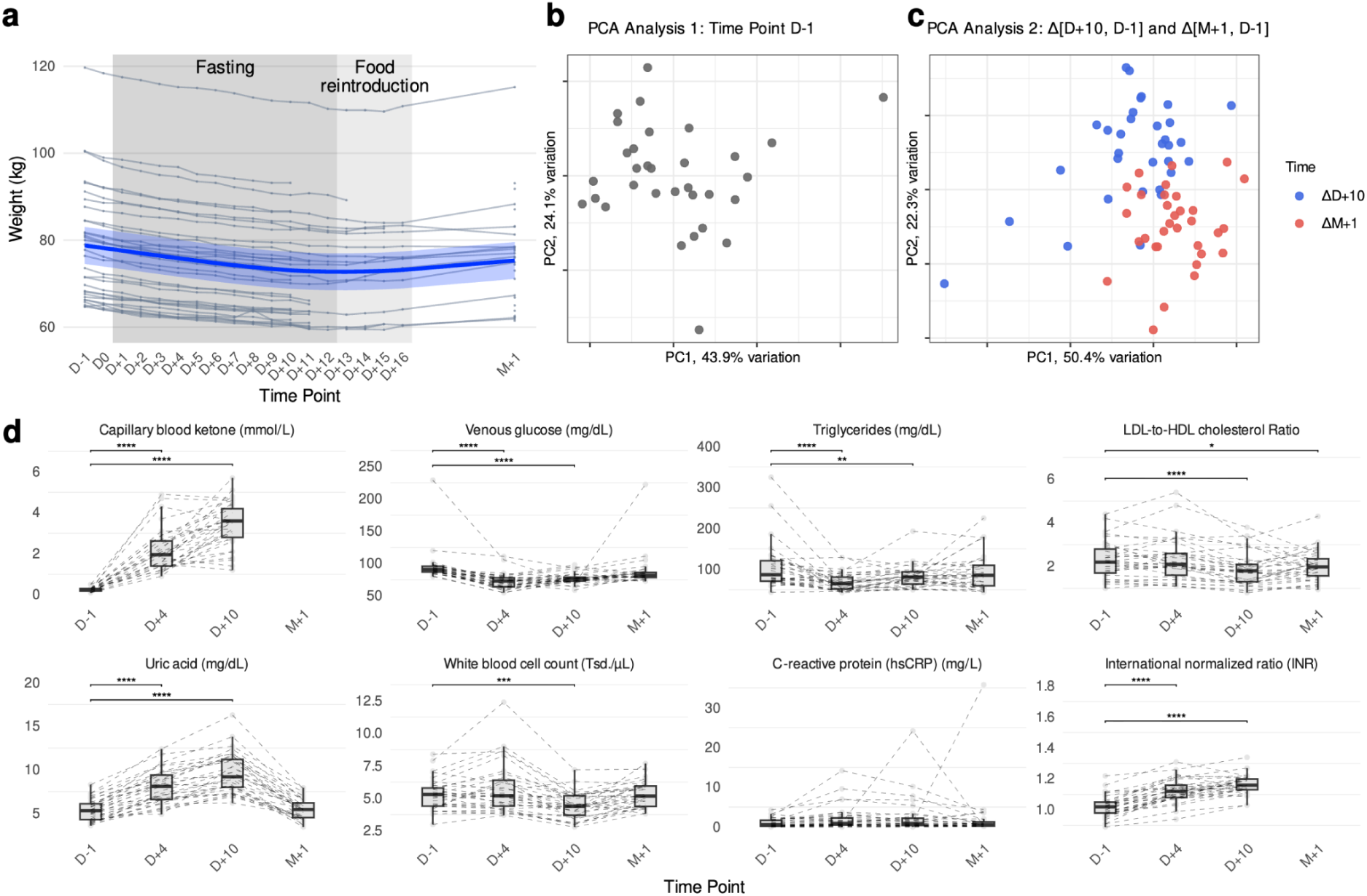
Systemic physiological changes during and after fasting. **a)** Daily body weight measurements collected from each participant from baseline (D-1) through the fasting and food reintroduction phases (D0 to D+16) and at the one-month follow-up (M+1). Shaded areas indicate fasting and food reintroduction periods; the blue bold line denotes the cohort mean. Weight measurements during later fasting and reintroduction time points were partially self-reported, resulting in increased missingness despite full participant retention. **b)** Principal component analysis (PCA) of standardized blood parameters measured at baseline (D−1), shown for the first two principal components. **c)** PCA of within-person changes in blood parameters relative to baseline, comparing D+10 vs D−1 and M+1 vs D−1; the first two principal components are shown. **d)** Selected clinical parameters plotted across study time points. Dashed lines indicate individual trajectories. Statistical significance for comparisons with baseline (D−1) is indicated as follows: * p < 0.05, ** p < 0.01, *** p < 0.001.

Body weight declined progressively during the fasting period, with an average reduction of approximately 6 kg by day 12 (-5.9 kg [5.2, 6.6], p<0.001), and increased partially during food reintroduction and at the one-month follow-up (-2.8 kg [0.4, 5.1], p = 0.015) (Figure 2a). Circulating ketone bodies increased during fasting, accompanied by reductions in circulating glucose and related indices of glycemic regulation, indicating a shift in energy metabolism away from glucose-based substrates toward lipid-derived fuels (Figure 2d; Table 2). In parallel, lipid profile measures—including triglycerides and the LDL/HDL ratio shown in Figure 2d, as well as additional lipid measures summarized in Table 2—changed during fasting and partially returned toward baseline values after resumption of habitual eating.

Together, these observations describe an acute metabolic response that was largely reversible following the intervention, with some parameters remaining shifted relative to baseline at one month.

To summarize multivariate changes across clinical parameters, principal component analysis (PCA) was performed on blood measurements at baseline (D−1) and separately on within-person change scores relative to baseline (Δ[D+10, D−1] and Δ[M+1, D−1]). The first two principal components (PCs) accounted for 77% of the variance at baseline and 71% for change scores, indicating a low-dimensional structure despite the number of measured parameters. Baseline PCs showed continuous inter-individual variation without evident clustering (Figure 2b). PCs of change scores showed separation by contrast, with Δ[D+10, D−1] and Δ[M+1, D−1] occupying different regions (Figure 2c). Additional PCs and PCA loadings for both analyses are shown in Supplementary Figure 2 and 3.

Beyond metabolic markers, fasting was accompanied by changes in parameters reflecting turnover, systemic stress, and regulatory processes (Figure 2d; Table 2). Uric acid levels increased during fasting and returned toward baseline at follow-up, indicating enhanced purine turnover under catabolic conditions. Leukocyte counts and inflammatory markers showed modest, time-dependent fluctuations; for example, hs-C-reactive protein slightly increased at D+10 and returned toward baseline at M+1, with high inter-individual variability and no statistically significant difference (Figure 2d; Table 2). Coagulation parameters (INR and PTT) shift substantially during fasting (Figure 2d; Table 2); although follow-up measurements at M+1 were not available, fasting-associated changes in coagulation markers have been reported to be transient in previous studies^51–53^.

Collectively, these results define the physiological context of the fasting intervention: a multivariate and largely reversible systemic response extending beyond metabolism to include hematological, inflammatory, and regulatory domains. The following analyses examine whether epigenetics capture transient fasting-associated states, potentially longer-lasting adaptations, or remain stable despite pronounced short-term physiological changes.

### DNA methylation landscape changes during and after fasting

*TSS200* denotes regions within 0–200 bp upstream of the transcription start site (TSS); *TSS1500* refers to regions 200–1500 bp upstream of the TSS; *1stExon* indicates the first coding exon of a gene, and *ExonBnd* refers to exon boundary regions.

To assess locus-specific DNA methylation changes associated with fasting, differential methylation analyses were performed comparing baseline (D−1) with day 10 of fasting (D+10) and with the one-month follow-up (M+1). CpG-wise tests were conducted using linear mixed-effects models adjusting for scan date, age, sex, and measured lymphocyte proportions (Methods). No differentially methylated CpG sites were detected between D−1 and D+10 at the applied significance threshold (Methods), indicating an absence of detectable CpG-level methylation changes during the acute fasting phase.

In contrast, comparisons between D−1 and M+1 identified subtle but widespread methylation shifts (Figure 3a). Both hypermethylated and hypomethylated sites were observed, with small to modest effect sizes. These changes were broadly distributed across genomic contexts, including CpG islands, shores, shelves, and open sea regions, as well as across gene-associated features such as promoter regions, gene bodies, and untranslated regions (Figure 3b,c). No prominent chromosomal clustering of differentially methylated CpGs was observed (Supplementary Figure 5).

**Figure 3:**
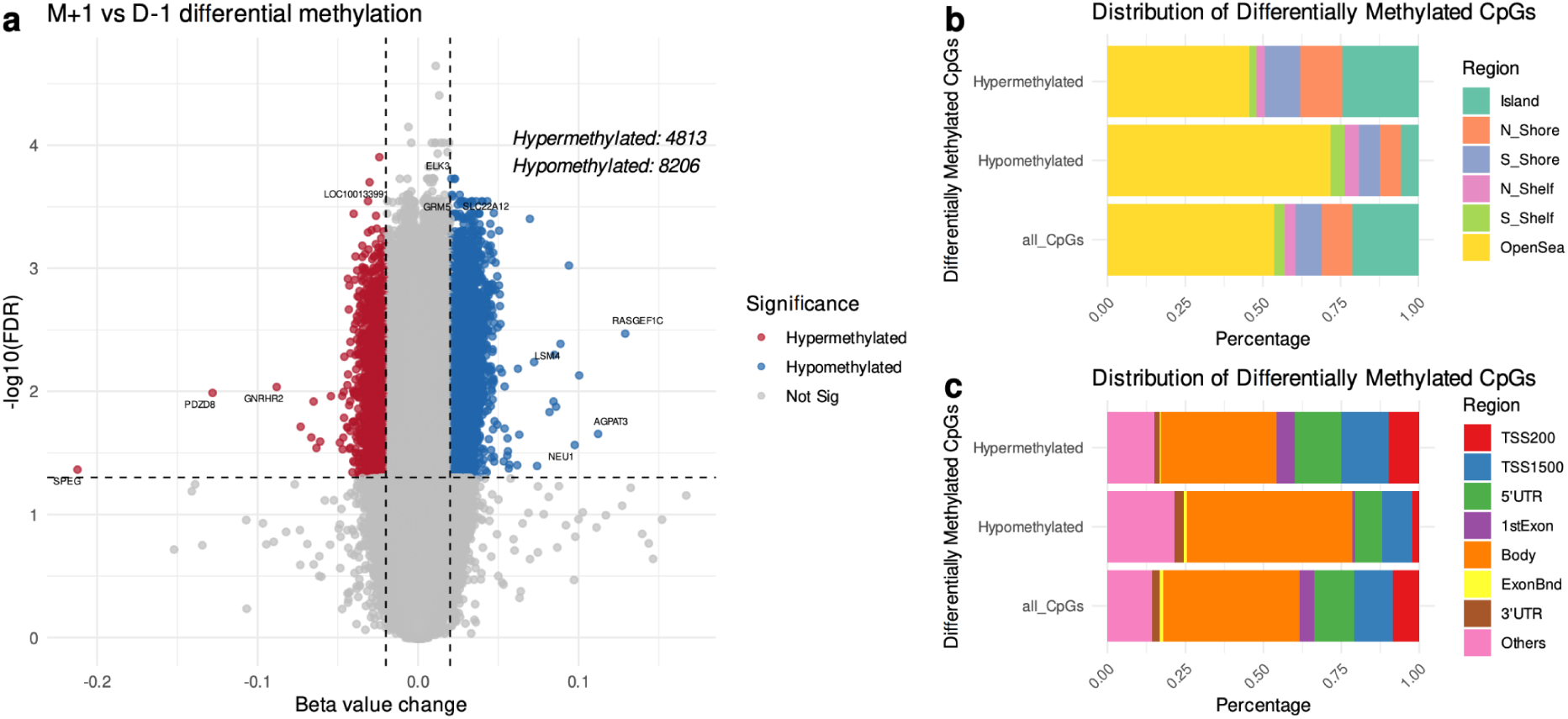
Differential DNA methylation analysis between baseline (D-1) and one-month follow-up (M+1). **a)** Volcano plot showing differentially methylated CpG sites, with top hits labeled by their associated genes. **b-c)** Genomic distribution of hyper- and hypo-methylated CpG sites compared to the distribution of all CpG probes on the array. CpGs are grouped by CpG island context (b) and by gene-associated features (c). Legend labels represent as follows: *island* refers to CpG islands; *N_shore* and *S_shore* indicate regions flanking CpG islands up to 2 kb upstream or downstream, while *N_shelf* and *S_shelf* represent regions 2–4 kb from CpG islands; *OpenSea* refers to isolated CpGs not in proximity to CpG islands.

Functional enrichment analyses based on Gene Ontology and KEGG pathways did not identify significant overrepresentation of biological processes or pathways among the differentially methylated CpGs. Together, these findings indicate that prolonged fasting was not associated with coordinated, pathway-level DNA methylation remodeling at individual CpG resolution in whole blood, but rather with sparse and dispersed methylation differences detectable at one month following the intervention, after accounting for major measured blood cell covariates.

### Dynamics of epigenetic aging during and after fasting

We assessed epigenetic aging using twelve established DNA methylation clocks spanning multiple clock generations and modeling paradigms. These included three first-generation chronological age clocks in their principal component (PC) versions (PCHorvath1, PCHorvath2, and PCHannum); second- and later-generation clocks capturing phenotypic aging, mortality risk, and pace of aging (PCPhenoAge, PCGrimAge, DNAmGrimAge2, DunedinPACE and SystemsAge); a PC-based DNA methylation estimator of telomere length (PCDNAmTL); and three causality-enriched clocks (AdaptAge, DamAge, and CausAge).

As expected, clocks explicitly trained to predict chronological age or age-associated outcomes—Horvath-, PhenoAge-, and GrimAge-based clocks—showed strong associations with chronological age across all samples and time points (marginal R^2^ = 0.83–0.94; Supplementary Figure 6). PC DNAmTL also exhibited clear age associations, consistent with prior reports. In contrast, DunedinPACE, which quantifies the pace of aging rather than accumulated age, showed minimal association with chronological age. SystemsAge and the causality-enriched clocks displayed age associations consistent with those reported in their original publications^27,31^.

To characterize fasting-associated changes in epigenetic aging, we quantified within-subject differences relative to baseline (D-1) at day 10 of fasting (D+10) and at the one-month follow-up (M+1). Changes were expressed as standardized differences in epigenetic age acceleration (ΔSRAA), scaled by the baseline standard deviation to facilitate comparison across clocks (Methods).

#### System-specific epigenetic remodeling during and after fasting

Epigenetic clocks exhibited heterogeneous temporal responses to fasting and food reintroduction (Figure 4a). At D+10, chronological-age trained clocks, including PCHorvath1, PCHorvath2, and PCHannum, showed modest increases relative to baseline, whereas DunedinPACE and PCDNAmTL decreased. PCGrimAge also showed a marginal increase during fasting. Changes varied substantially between individuals. At the one-month follow-up (M+1), PCHorvath2 remained elevated, while SystemsAge and DamAge showed delayed increases during recovery. In contrast, AdaptAge decreased at M+1, whereas several other clocks partially returned toward baseline. Despite divergent temporal trajectories across individual clocks, responses occupied a relatively low-dimensional space, with related clock families exhibiting partially shared loading patterns (Supplementary Figure 14).

**Figure 4:**
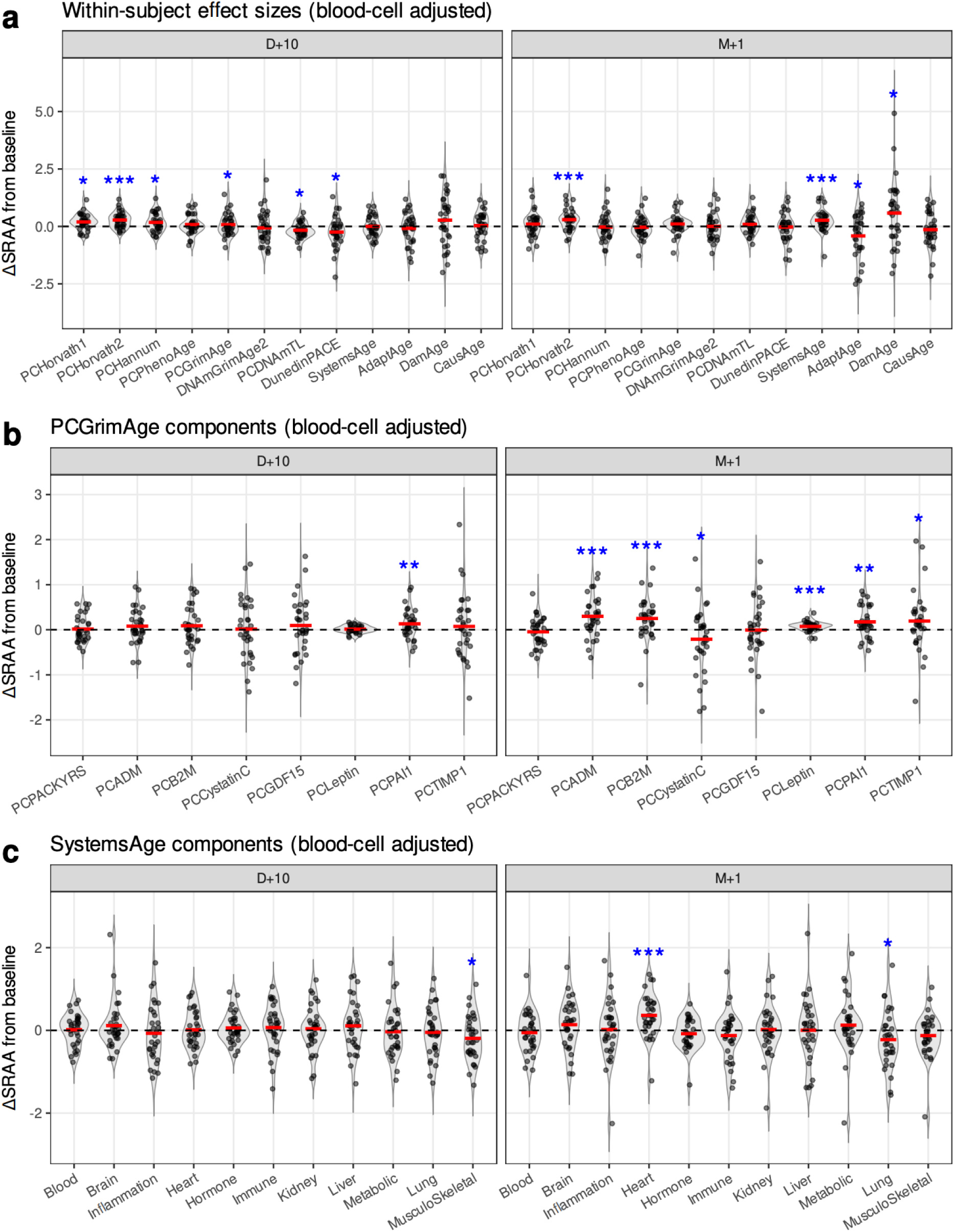
Responses of epigenetic clocks during and after fasting. Within-subject changes from baseline in epigenetic age acceleration (ΔSRAA) at day 10 of fasting (D+10) and one month after food reintroduction (M+1), expressed in units of baseline standard deviation, are shown for **a)** all 12 clocks, **b)** PCGrimAge components, and **c)** SystemsAge components. Each dot represents an individual participant; red horizontal bars indicate the mean change across individuals; the dashed line marks no change relative to baseline. Blue asterisks indicate statistically significant contrasts versus baseline derived from linear mixed-effects models, with fixed-effect covariates including sex **and lymphocyte proportions**, and subject modeled as a random effect (*P < 0.05, **P < 0.01, ***P < 0.001).

Component-level analyses of DNAm GrimAge and SystemsAge revealed heterogeneous trajectories across physiological systems and biomarker surrogates (Figure 4b–c). Component analyses were primarily performed using the PC-based implementation because surrogate trajectories differed substantially across GrimAge implementations (Supplementary information Section 4). Within DNAm GrimAge components, DNAm Plasminogen Activation Inhibitor 1 (PAI-1), a surrogate associated with fibrinolysis, metabolic syndrome, triglycerides, visceral adiposity, and fatty liver biology, increased already during fasting at D+10 and remained elevated at M+1. DNAm Adrenomedullin (ADM) (Figure 4c), linked to cardiovascular and vascular regulation, and DNAm Beta-2 Microglobulin (B2M), associated with systemic inflammation, also increased at M+1. In contrast, the DNAm Cystatin C surrogate, a marker associated with kidney dysfunction and metabolic disease risk, decreased during recovery (Figure 4b). The leptin surrogate, reflecting adipose tissue state and systemic energy balance and previously reported to be inversely associated with mortality risk, increased at M+1.

SystemsAge components also exhibited divergent system-specific trajectories (Figure 4c). The Heart score increased during recovery at M+1, whereas the Musculoskeletal score, which has been associated with physical function, frailty, diabetes, and metabolic health, decreased during fasting at D+10 (Figure 4c).

Together, these findings suggest heterogeneous remodeling across vascular, metabolic, inflammatory, musculoskeletal, and endocrine-associated methylation signatures during fasting and food reintroduction, rather than a uniform shift across biological aging measures.

These patterns were largely robust to lymphocyte adjustment, with adjusted and unadjusted effect estimates showing strong concordance across clocks (Supplementary Figure 7, 10, and 12). Decomposition analyses further indicated that fasting-associated clock changes were predominantly captured by residual, composition-independent components rather than lymphocyte-associated variation (Supplementary Figure 9, 11, and 13). See Supplementary Information for more details.

### Epigenetic aging and individual responsiveness to fasting

We next examined whether epigenetic aging measures were associated with inter-individual variability in fasting responsiveness, quantified as maximum percentage weight loss during the intervention. Epigenetic age acceleration was defined as residuals from regressing clock-predicted age on chronological age. Associations were tested using linear mixed-effects models (Methods).

Across multiple clocks, greater weight loss was consistently associated with lower epigenetic age acceleration (Figure 5; Supplementary Figure 18). Horvath-based age acceleration showed a nominally significant negative association with maximum weight loss (β = −94.4 to −130 across PC Horvath clocks; p = 0.038 and p = 0.0066, respectively). PhenoAge acceleration showed a similar association (β = −143, p = 0.0258). DunedinPACE also exhibited a negative association with weight loss (β = −2.0, p = 0.0504). Other clocks, including GrimAge, GrimAge v2, SystemsAge, and CausAge, showed directionally consistent negative coefficients, although these associations did not reach statistical significance.

**Figure 5:**
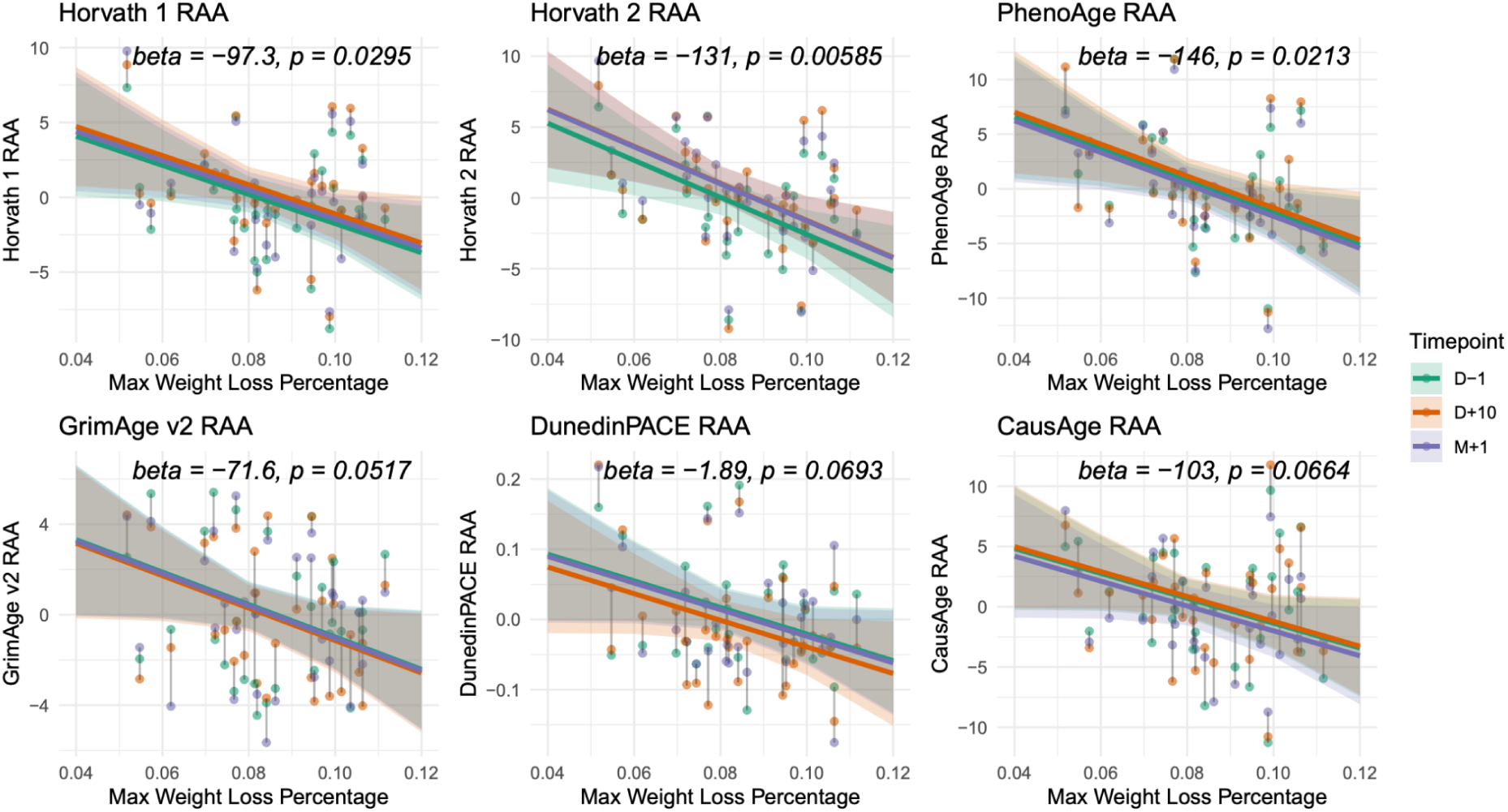
Association between residual epigenetic age acceleration and maximum weight loss percentage for 6 representative clocks. Residual age acceleration was modeled using linear mixed-effects models including maximum weight loss percentage, time point, sex, and lymphocyte proportions as fixed effects, with participant ID as a random effect. The marginal effect of maximum weight loss percentage at each time point is shown as a colored regression line, with the corresponding regression coefficient (beta) and p-value (p) displayed in each panel. Data points are colored by time points and connected by lines to indicate repeated measures from the same individual. Corresponding analyses using the same model specification are shown for all clocks in Supplementary Figure 18. Results from models additionally adjusted for baseline BMI are shown in Supplementary Figure 19.

Adjustment for baseline BMI did not materially alter the effect estimates (Supplementary Figure 19), although nominal significance was slightly reduced, consistent with increased estimation uncertainty rather than substantial BMI confounding. These results suggest that the observed associations are not primarily driven by baseline adiposity. In addition, we found no evidence for interaction between BMI and weight loss in predicting epigenetic age acceleration (Methods, Supplementary Table 2), indicating that the association between fasting responsiveness and epigenetic aging measures does not substantially differ across BMI levels. Baseline-only sensitivity analyses yielded directionally consistent associations across clocks, although with reduced statistical significance compared to repeated-measures models (Methods, Supplementary Table 3).

Together, these results indicate that individuals with younger-appearing epigenetic profiles or slower aging pace tended to exhibit greater physiological responsiveness to prolonged fasting. While causal inference is not possible, this pattern is consistent with the hypothesis that epigenetic aging measures capture aspects of baseline biological state related to metabolic adaptability under energetic stress. The modest effect sizes, partial significance across clocks, and slight increases in estimation uncertainty following BMI adjustment underscore the need for larger cohorts to determine the robustness and specificity of these associations.

## Discussion

In this longitudinal interventional study, we examined the physiological and epigenetic effects of a medically supervised 12-day prolonged fasting regimen in humans. Fasting induced pronounced systemic metabolic adaptations, some of which remained shifted at one month, whereas CpG-level DNA methylation changes were sparse, modest in effect size, and dispersed, with no evidence of coordinated pathway-level remodeling. In contrast, multiple epigenetic aging measures showed clock-specific, time-dependent responses during fasting and after food reintroduction, with substantial inter-individual heterogeneity and structured relationships across clocks. Together, these findings indicate a dissociation between acute physiological adaptation, locus-specific DNA methylation changes, and integrative epigenetic aging signals during and after prolonged fasting.

The physiological changes observed—including the metabolic switch toward lipid-derived substrates, altered glycemic regulation, lipid profiles, and markers of catabolic stress—are consistent with previous clinical and experimental studies^5,15,54^ and reflect responses to an acute energy deficit. Most markers returned toward baseline after food reintroduction, indicating state-dependent metabolic adaptation rather than durable modifications of long-term cardiometabolic risk. Conventional clinical biomarkers are optimized for steady-state risk stratification. Epigenetic aging measures, by contrast, offer a complementary, DNA-based readout that integrates distributed molecular responses and may capture broader regulatory and cellular states operating over longer timescales.

This divergence between acute physiology and epigenetic signals is evident at two epigenetic levels. At the level of individual CpG sites, the absence of coordinated DNA methylation changes, particularly during fasting, likely reflects differences in biological timescales and regulation levels. Acute metabolic perturbations primarily involve rapid shifts in substrate utilization, hormone signaling, and enzymatic activity, whereas stable CpG-level DNA methylation changes may require longer or repeated exposures, cell turnover, or sustained changes in cellular composition^55,56^. In whole blood, modest locus-specific methylation effects may also be diluted by cellular heterogeneity despite adjustment for measured immune cell proportions^57,58^. Thus, the limited CpG-level signals indicate that time-limited fasting does not induce gene-centric methylation remodeling detectable at individual CpG resolution, at least within the temporal window examined.

In contrast, epigenetic aging clocks, which aggregate information across hundreds of CpGs^24,26,29,30^, showed measurable, clock-specific responses. During fasting, several chronological-age clocks and one mortality-associated clock increased and partially rebounded thereafter, consistent with reports that some biological age metrics transiently rise under acute physiological stress^59^. In contrast, the pace-of-aging measure DunedinPACE^28^ decreased during fasting. Unlike clocks trained to estimate accumulated biological age, DunedinPACE captures the current rate of physiological decline, which may be more sensitive to transient metabolic slowing during fasting without necessarily requiring reversed accumulated epigenetic aging.

Component-level analyses further revealed that composite epigenetic aging measures aggregate physiologically distinct responses across biological systems that do not necessarily shift uniformly toward either a biologically “older” or “younger” state. At D+10 and M+1, the increase in DNAm PAI-1 may reflect metabolic and hepatic remodeling associated with large shifts in energy balance and lipid mobilization during and after fasting. Similarly, increases in DNAm ADM and the SystemsAge Heart score at M+1 may reflect vascular and cardiovascular remodeling processes during recovery.

In contrast, the decreases in SystemsAge Musculoskeletal score and DNAm Cystatin C during fasting or recovery may reflect improvements in metabolic health states linked to insulin sensitivity and systemic metabolic burden. The leptin surrogate increased at M+1, potentially consistent with the known suppression of leptin during prolonged fasting^60^ and subsequent restoration of adiposity-associated signaling during recovery.

Together, these component-level patterns suggest that transient increases or decreases in some “risk-associated” biomarkers may not necessarily indicate accelerated or rejuvenated biological aging, but instead reflect coordinated physiological adaptation related to energetic stress, lipid mobilization, and vascular adaptation. These findings further highlight that composite clock outputs can obscure biologically distinct underlying trajectories, particularly during acute interventions or under non-equilibrium physiological states.

Our findings are consistent with prior dietary intervention studies^38^, including the CALERIE trial^33^, a two-year caloric restriction study in humans. Analyses of epigenetic aging outcomes in CALERIE reported modest average effect sizes, substantial inter-individual variability, and limited consistency across different epigenetic aging measures, with only DunedinPACE showing a consistent decrease^33^. Together with our findings, this suggests that DunedinPACE may be particularly well suited for detecting beneficial physiological responses to caloric restriction and fasting interventions. Furthermore, exploratory principal component analyses supported the presence of structured, partially shared response patterns across clocks, rather than purely idiosyncratic clock behavior. These multivariate clock response dimensions were largely not explained by routine circulating metabolic markers, suggesting that epigenetic aging measures do not simply recapitulate acute metabolic flux. Notably, one lower-variance component dominated by CausalAge remained associated with insulin resistance, consistent with prior analyses of CausalAge showing enrichment of mediator genes in metabolic and nutrient-sensing pathways, including fructose and mannose metabolism, cholesterol and sterol catabolism, and mTOR signaling^31^.

Our findings also align with broader evidence suggesting that adaptive physiological remodeling and aging-related decline are not necessarily equivalent biological processes. In genetically diverse mice, lifespan extension from dietary restriction is not predicted by classic metabolic traits like adiposity or fasting glucose, but by factors like stress resilience, immune function, and genetic background^61^. Similarly, studies of naturally recurring fasting states such as hibernation show that acute molecular stress during energy deficit (e.g., telomere attrition) can coexist with slower organismal aging over repeated cycles^62,63^. Together, these observations frame our human data within a broader principle: short-term molecular responses to fasting primarily capture adaptive state transitions, while aging-related adaptations operate on longer, possibly cumulative, timescales.

Furthermore, baseline epigenetic age acceleration was associated with variability in fasting responsiveness, suggesting these measures may capture aspects of metabolic adaptive capacity. Aging has been conceptualized not only as the accumulation of molecular damage, but also as a progressive decline in the ability to maintain or restore homeostasis following physiological stressors^64^. In this context, differential epigenetic aging states may influence the capacity to adapt to energetic stress and metabolic switching during prolonged fasting, consistent with broader concepts of metabolic flexibility and systemic resilience^65^. However, because these measurements were derived from peripheral blood rather than metabolically active tissues, the underlying mechanisms remain unresolved.

This study has several limitations. The sample size was modest, limiting power to detect small effects and to resolve clock-specific differences with high resolution. The study used a single-arm longitudinal design without a parallel control group; although within-subject comparisons reduce inter-individual variability, the absence of a control group limits the ability to fully disentangle fasting effects from other time-dependent factors. DNA methylation was measured in whole blood, which may obscure cell-type–specific epigenetic changes despite adjustment for measured immune cell proportions^57,58^. The follow-up period was limited to one month, precluding assessment of longer-term persistence or cumulative effects of repeated fasting cycles. Finally, although epigenetic clocks are established predictors of disease and mortality, their validity as surrogate endpoints for short-term interventions remains incompletely characterized^38^.

Future studies should incorporate longer follow-up, repeated interventions, cell-type resolved epigenetic profiling—particularly in metabolically relevant tissues—and integration with other molecular layers such as transcriptomics or chromatin accessibility. Such approaches will be essential to clarify how fasting-induced physiological adaptations interface with epigenetic regulation and aging-related processes across biologically meaningful timescales.

## Author contributions

S.L. contributed to formal analysis, software development, data curation, investigation, visualization, writing of the original draft and manuscript revision. M.K. contributed to data curation, investigation, visualization, manuscript drafting, and manuscript revision. F.v.M. contributed to project administration, resources, supervision, investigation, and manuscript revision. R.M. contributed to study conceptualization, project administration, resources, supervision, investigation, manuscript drafting, and manuscript revision. C.K., L.C., S.N., and S.G.N. contributed to investigation and manuscript revision. S.H. contributed to manuscript revision. F.G. contributed to study conceptualization, project administration, investigation, and manuscript revision. F.W.d.T. contributed to study conceptualization, supervision, and manuscript revision.

## Author information

Correspondence should be addressed to Robin Mesnage and Ferdinand von Meyenn.

## Funding

F.v.M. acknowledges funding from ETH Zurich (core funding) and the European Research Council (StG #803491, and AdG #101200612). R.M. acknowledges funding from Buchinger Wilhelmi Development & Holding GmbH.

## Ethics approval

The complete study protocol, including sample size considerations and clinical procedures, is described in the GENESIS protocol (lonG tErm fastiNg multi-systEm adaptationS In humanS; 10.3389/fnut.2022.951000^16^). The study was approved by the Ethics Committee of the Baden-Württemberg Medical Council (F-2021-075, 26 July 2021) and registered at ClinicalTrials.gov (NCT05031598). All participants provided written informed consent prior to enrollment.

## Acknowledgments

We wish to acknowledge all members of the von Meyenn Lab and Mark D. Robinson for helpful discussion and support.

## Conflicts of interest

Authors M.K.. F.G., F.W.T. and R.M. are employees of the Buchinger Wilhelmi

Development and Holding GmbH, Überlingen. CK, LC, SN, SGN are employees of GenKnowme. The Regents of the University of California are the sole owner of patents and patent applications directed at epigenetic biomarkers for which Steve Horvath is a named inventor. S.H. is a founder and paid consultant of the non-profit Epigenetic Clock Development Foundation that licenses these patents. The remaining authors declare that the research was conducted in the absence of any commercial or financial relationships that could be construed as a potential conflict of interest.

## Data availability

The datasets will be made available upon publication of the manuscript.

## Code availability

Notebooks, R scripts, and supporting data used for processing datasets and generating all the visualizations in this manuscript are available on GitHub: https://github.com/RoseYuan/fasting_paper.

## Supplementary information

### 1. Sensitivity of clock time effects to blood cell composition

Fasting can induce transient changes in white blood cell composition, which may contribute to bulk DNA methylation clock signals. To avoid over-correcting potentially relevant biological variation, we additionally performed time-effect analyses without lymphocyte adjustment and compared time effects estimated from models with and without lymphocyte adjustment (Supplementary Figure 7, 10, and 12). Across clocks, estimated effect sizes were highly similar before and after adjustment, with most estimates lying close to the identity line, indicating that lymphocyte composition was not the dominant driver of fasting-associated clock responses.

Nevertheless, systematic timepoint-dependent patterns were observed. At D+10, lymphocyte adjustment modestly increased estimated effect sizes for several clocks, indicated by the warm colours, whereas at M+1 adjustment more frequently attenuated the observed effects, indicated by the cold colours. Examination of within-subject lymphocyte changes revealed substantial inter-individual variability at both time points (Supplementary Figure 8). However, the coefficient of variation of lymphocyte changes was markedly higher at D+10 than at M+1 (2330% vs 831%), indicating that early fasting responses were substantially more heterogeneous across individuals. These observations suggest that acute fasting induces heterogeneous immune redistribution that contributes primarily to inter-individual variability at D+10, partially masking underlying clock changes, whereas later immune adaptation contributes more coherently to the observed group-level signal.

To further dissect these effects, we decomposed each clock signal into lymphocyte-associated and residual components (Supplementary Figure 9, 11, and 13). Across clocks, significant fasting-associated time effects were consistently detected in the residual component rather than the lymphocyte-associated component, indicating that the dominant clock changes were largely independent of blood cell composition. Together, these analyses suggest that lymphocyte composition influences the detectability of fasting-associated clock changes but does not constitute the primary driver of the observed epigenetic dynamics.

### 2. Multivariate structure of epigenetic aging responses

To examine shared structure across clock responses, we performed PCA on within-subject ΔSRAA at D+10 and M+1 (Supplementary Figure 14b). At both timepoints, the response space was low-dimensional: the first two principal components explained 42.4% and 21.0% of variance at D+10 and 49.1% and 23.3% at M+1, respectively.

Clock loadings on the dominant principal components revealed structured relationships consistent with clock training paradigms. Chronological-age–trained clocks grouped more closely together, as did health/risk-trained clocks, consistent with prior large-scale comparisons and intervention response benchmarking studies^54,66^. ￼Pace and telomere-related measures occupied distinct regions.

Causality-enriched clocks dominated a separate component (PC3), indicating an additional response dimension that was partially orthogonal to the age- and risk-related axes.

The PCA-derived structure was robust to sensitivity analyses excluding high-variance clocks, with dominant components preserved in variance explained, loading patterns, and sign, indicating that the observed low-dimensional structure reflects shared cross-clock response patterns rather than dominance by individual measures.

Together, these results provide additional confidence that the heterogeneous clock trajectories observed in the main analyses reflect structured, reproducible response patterns rather than random or clock-specific noise.

### 3. Relationship to clinical and metabolic measures

Participant scores along the first two PCA axes were not significantly associated with routine clinical or metabolic parameters, including weight loss, ketone levels, glucose, insulin, lipid measures, or hematological markers after correction for multiple testing (Methods; Supplementary Table 4). In contrast, a lower-variance component (PC3) at D+10—dominated by the causality-enriched clocks DamAge, AdaptAge, and CausAge—showed significant associations with insulin resistance-related measures, including HOMA index (ρ = 0.58, FDR = 0.005), insulin (ρ = 0.55, FDR = 0.006), and glucose (ρ = 0.51, FDR = 0.009). Consistently, changes in DamAge and AdaptAge at D+10 were individually associated with contemporaneous insulin resistance measures (Supplementary Figure 158-169).

Together, these findings suggest that the dominant axes of epigenetic clock response during fasting are largely independent of conventional clinical markers, whereas secondary response dimensions retain sensitivity to metabolic state.

### 4. Sensitivity of GrimAge component trajectories to clock implementation

To assess the robustness and implementation dependence of surrogate component trajectories, we additionally evaluated the corresponding components from the original GrimAge and GrimAge2 models. Several component trajectories differed substantially across implementations, including directionally inconsistent changes for certain surrogates such as DNAmADM (Supplementary Figure 17). These differences likely reflect the distinct statistical properties of PC-transformed versus original clock implementations, which have previously been shown to differ in technical reliability and longitudinal behavior. Given the improved robustness of PC-based clocks in longitudinal settings^32^, main-text component analyses focused on the PCGrimAge implementation.

## Supplementary figures

**Supplementary Figure 1:**
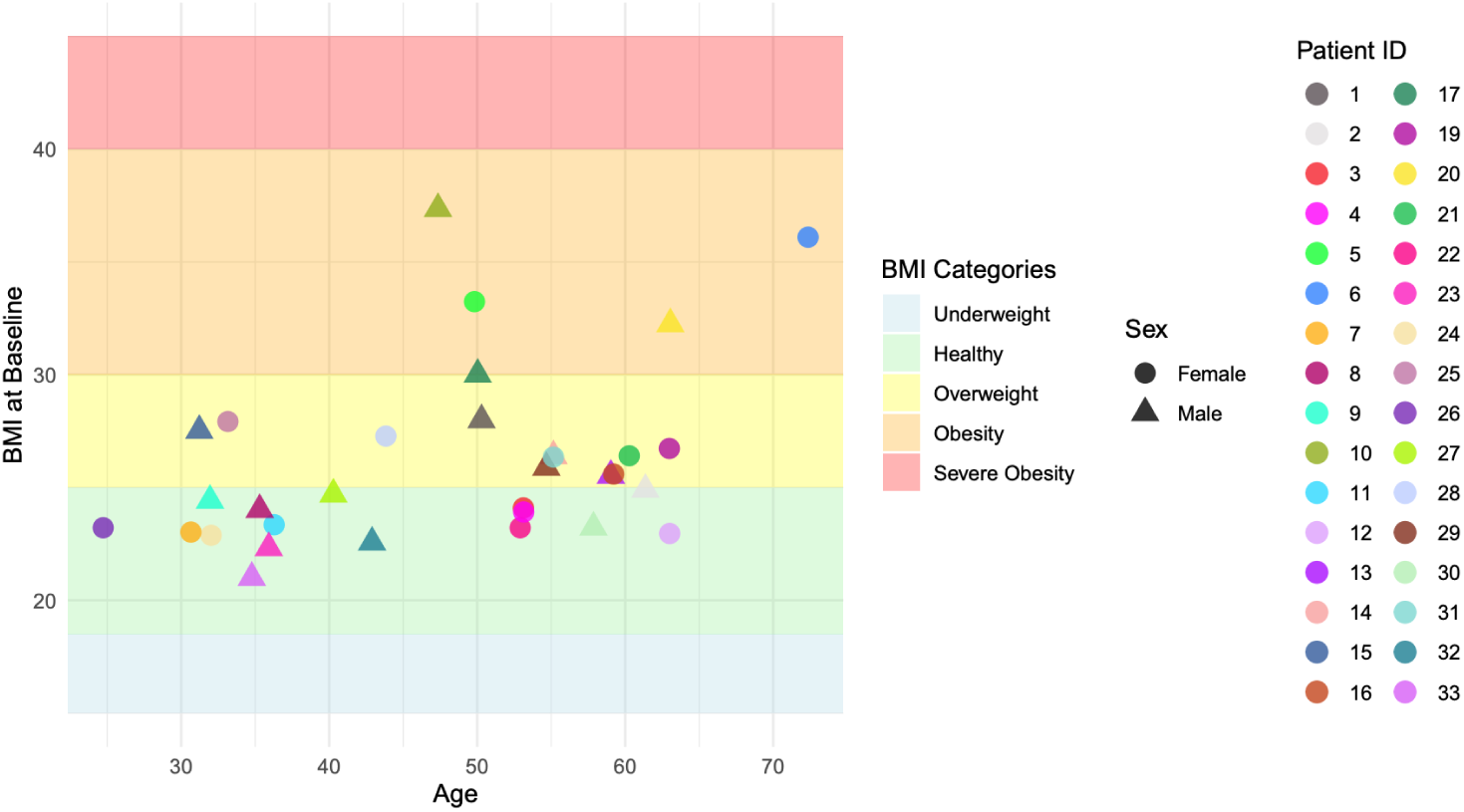
Additional clinical parameters. Chronological age, sex, and body mass index (BMI) measured at baseline (D-1).

**Supplementary Figure 2:**
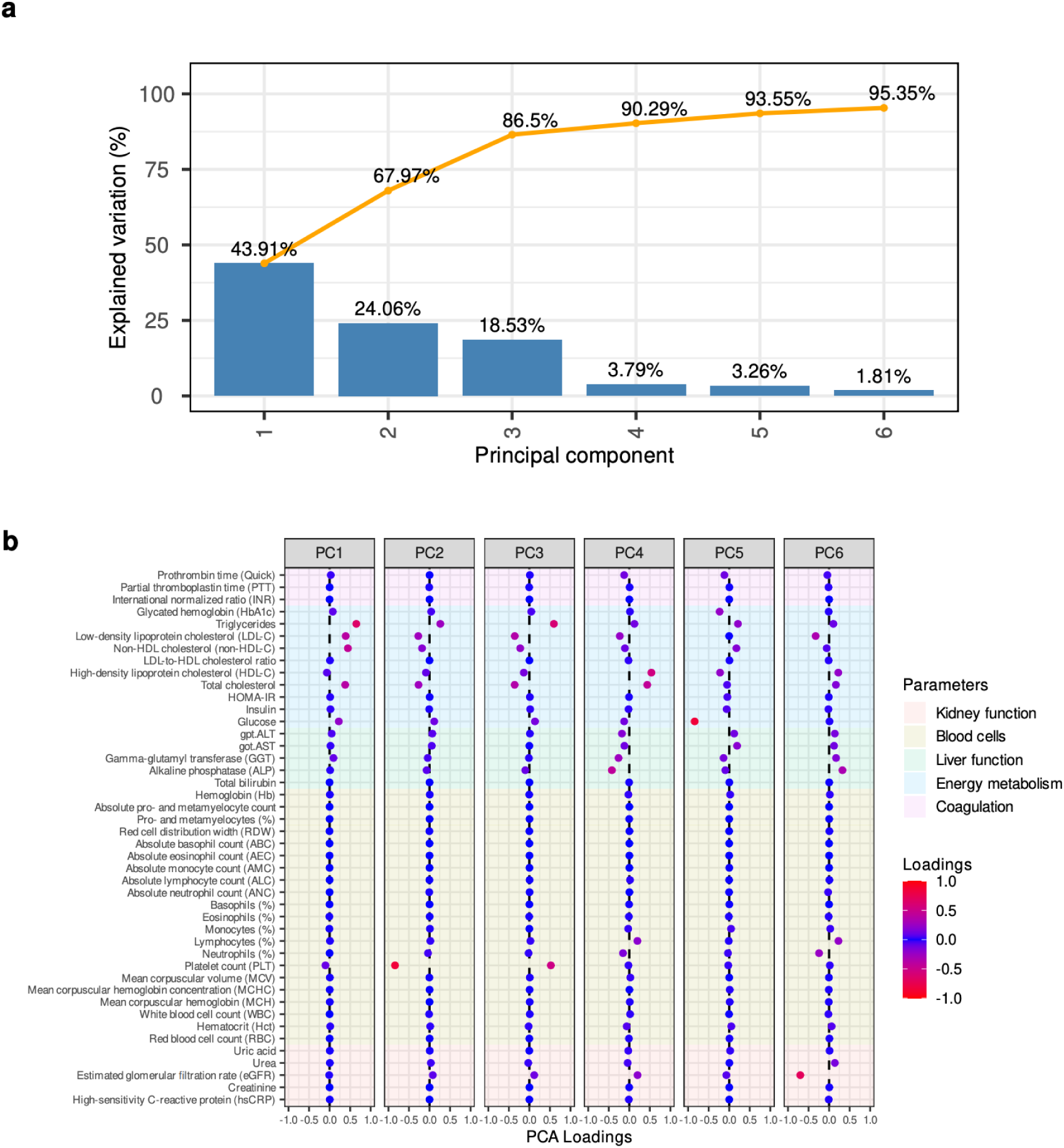
PCA of baseline measurements (D−1) shown in. Figure 2b**. a)** Scree plot showing the proportion of variance explained by the first 6 PCs. **b)** Loadings of individual blood parameters for the first 6 PCs.

**Supplementary Figure 3:**
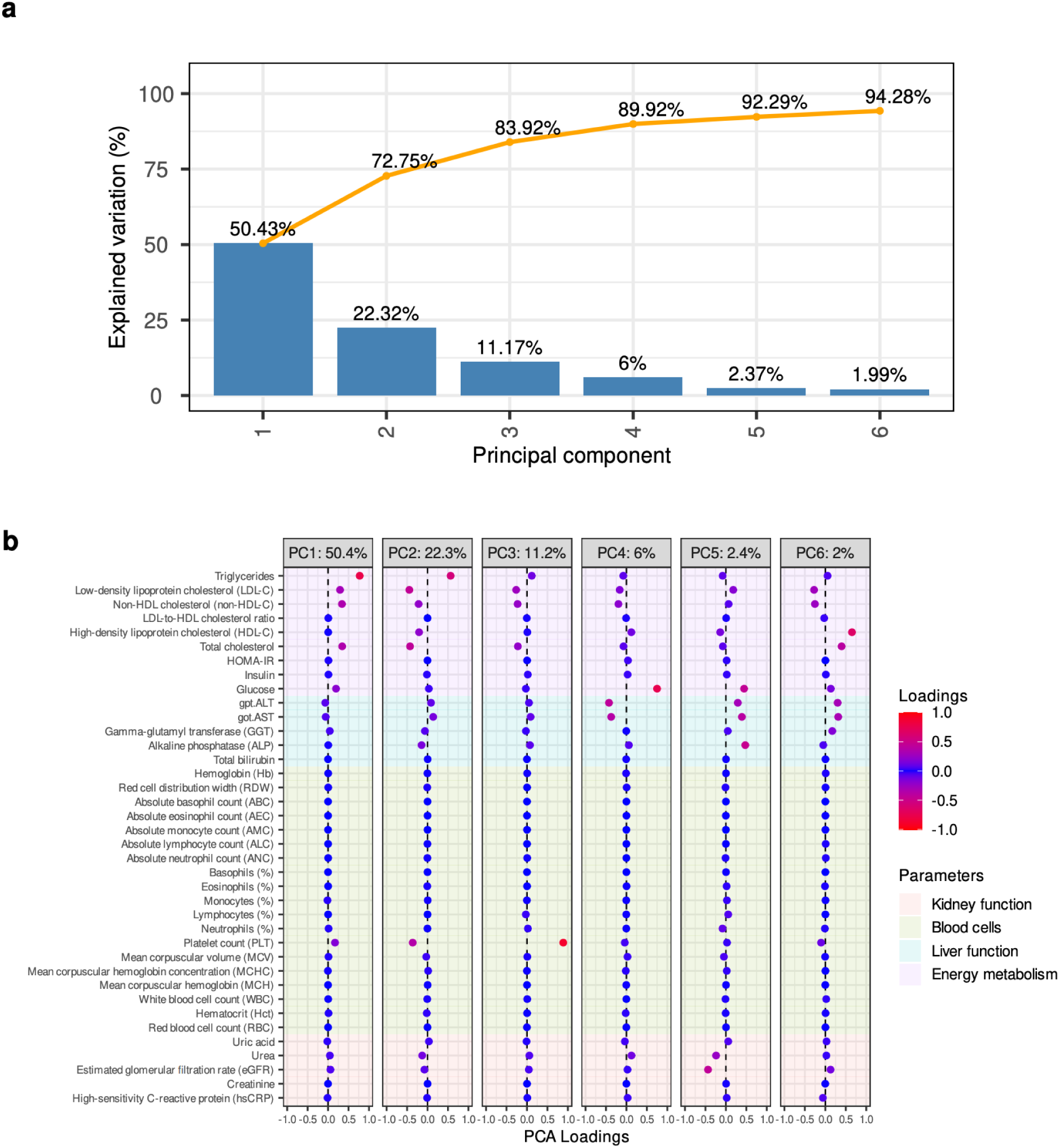
PCA of within-person changes as shown in Figure 2c. **a)** Scree plot showing the proportion of variance explained by the first 6 PCs. **b)**Loadings of within-person change scores for individual blood parameters.

**Supplementary Figure 4:**
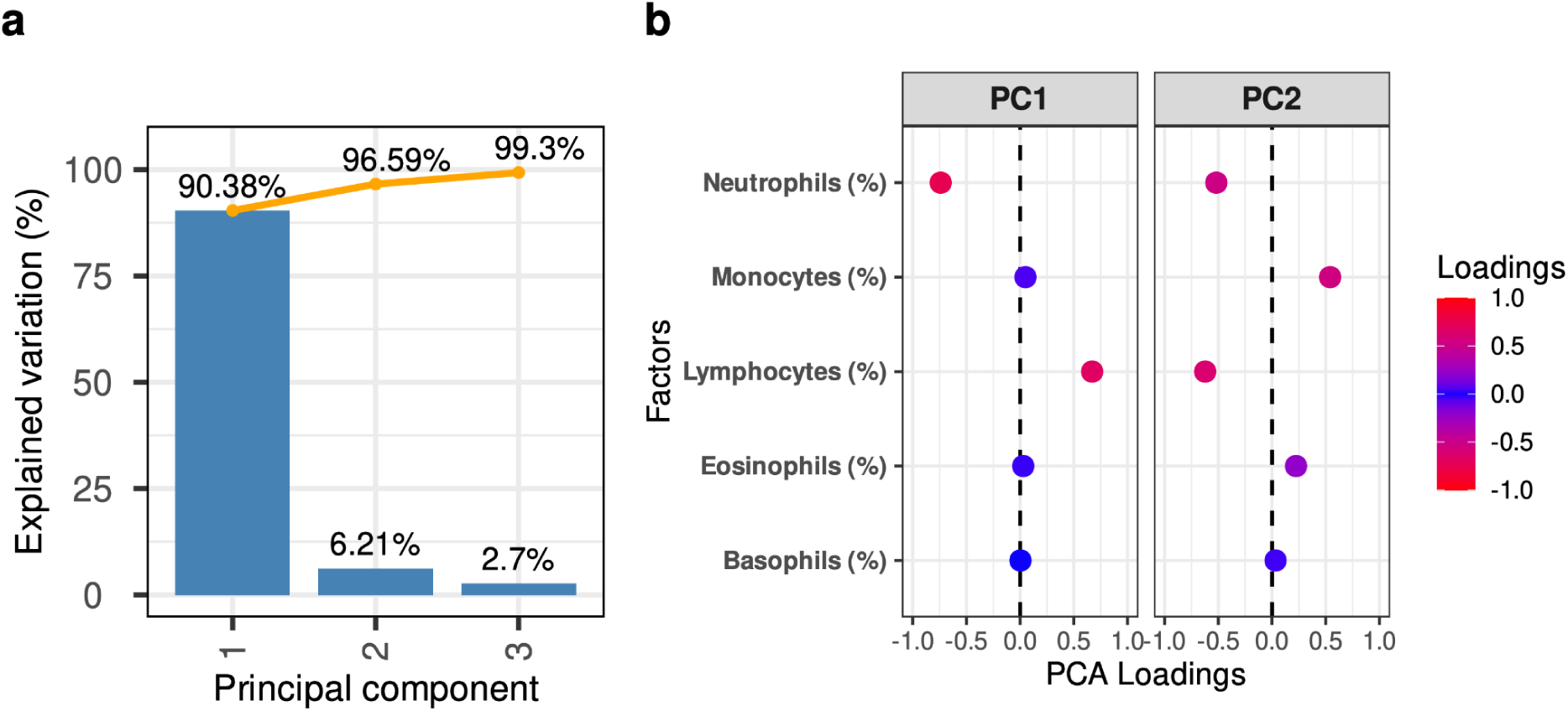
PCA of white blood cell composition measures. **a)** Scree plot showing the proportion of variance explained by the first 3 PCs. **b)** Loadings of individual white blood cell composition parameters for the first two PCs.

**Supplementary Figure 5:**
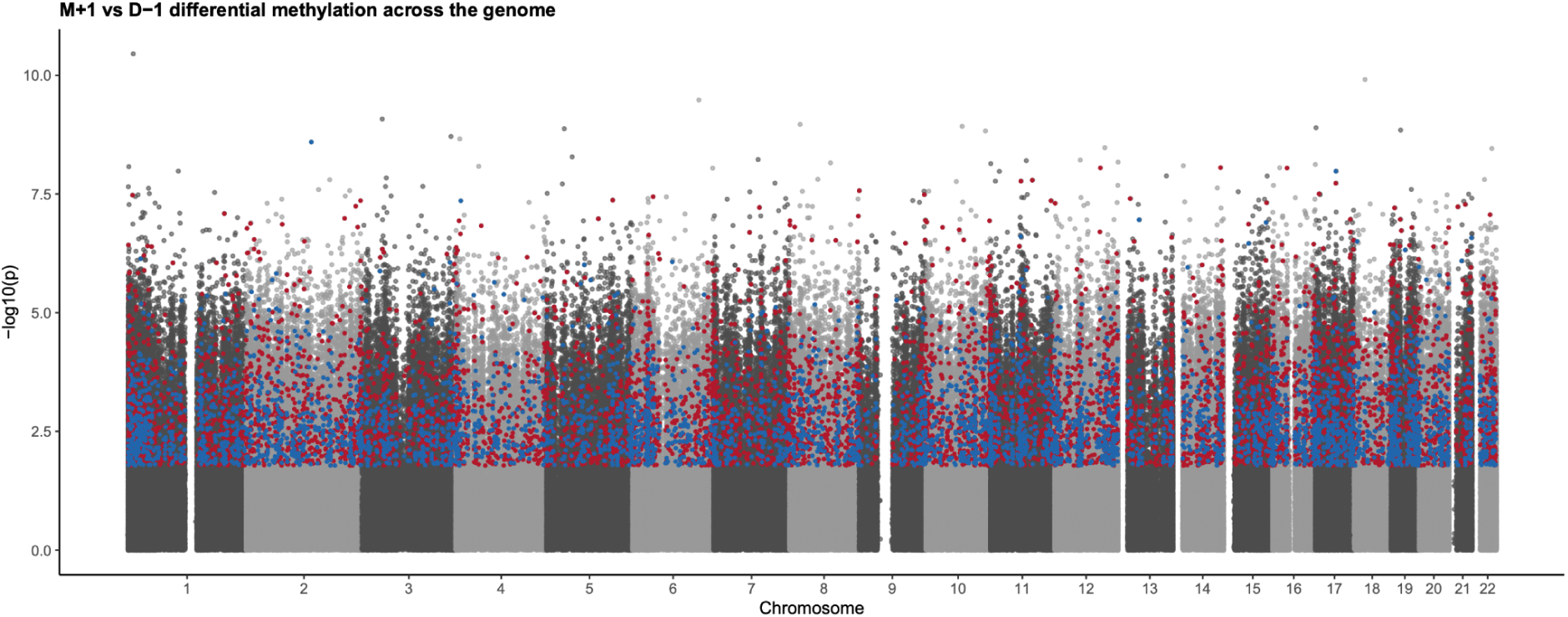
Differential DNA methylation at one-month follow-up. Manhattan plot showing differentially methylated CpG sites comparing one-month follow-up (M+1) to baseline (D−1). Each point represents a CpG site on the Illumina EPIC array, plotted by genomic position. Hypermethylated CpGs are shown in red and hypomethylated CpGs in blue, defined by an adjusted p-value < 0.05 and an absolute change in beta value > 0.02.

**Supplementary Figure 6:**
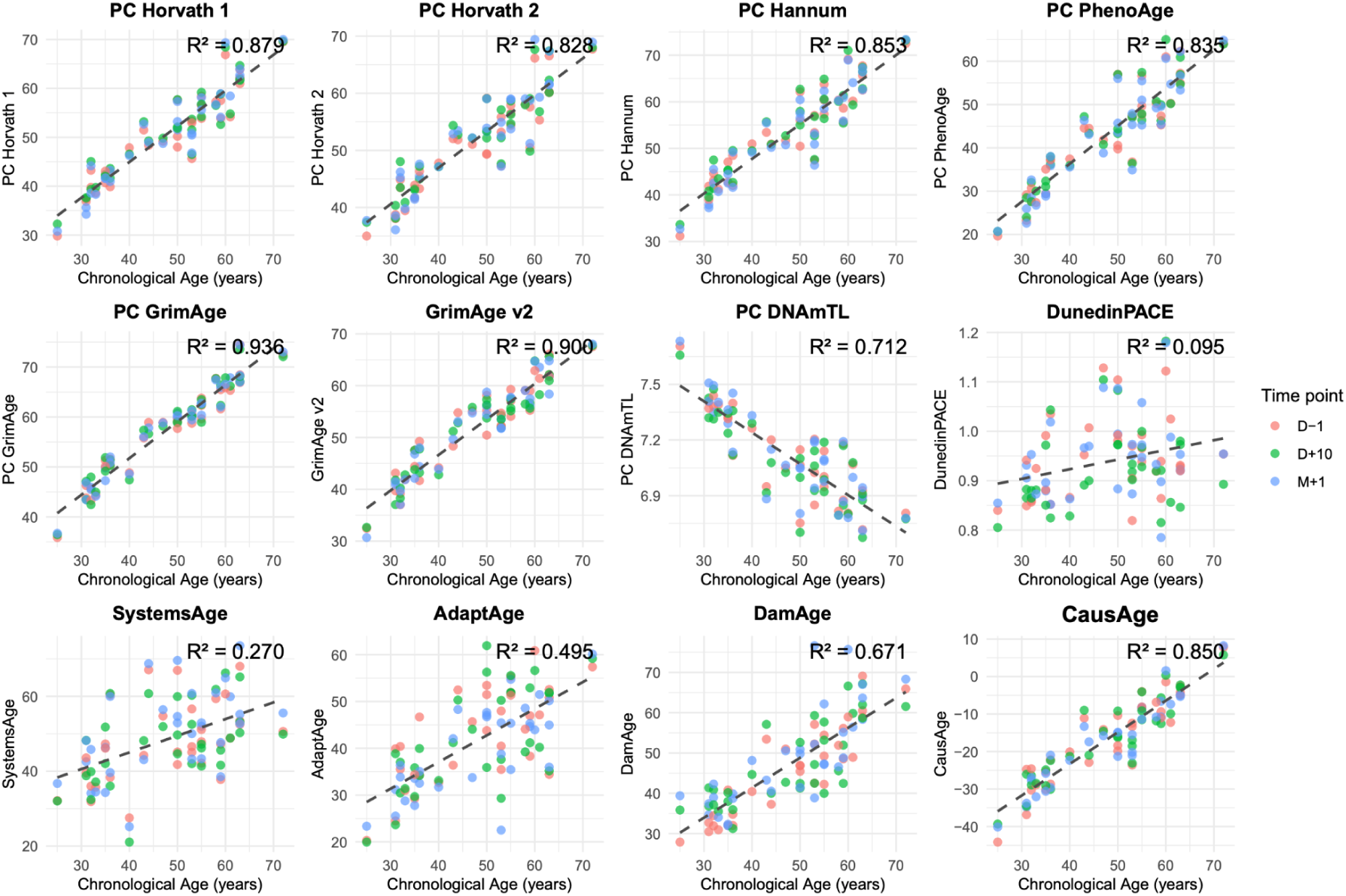
Association between epigenetic clock estimates and chronological age. Epigenetic age estimates are plotted against chronological age across all samples and time points (D−1, D+10, and M+1). Points are colored by time points. Dashed lines represent fixed-effects predictions from linear mixed-effects models with participant ID specified as a random effect and chronological age and time point included as fixed effects, averaged across time points. Reported R^2^ values correspond to the marginal R^2^ from the mixed-effects models, reflecting the proportion of variance in epigenetic age explained by chronological age.

**Supplementary Figure 7:**
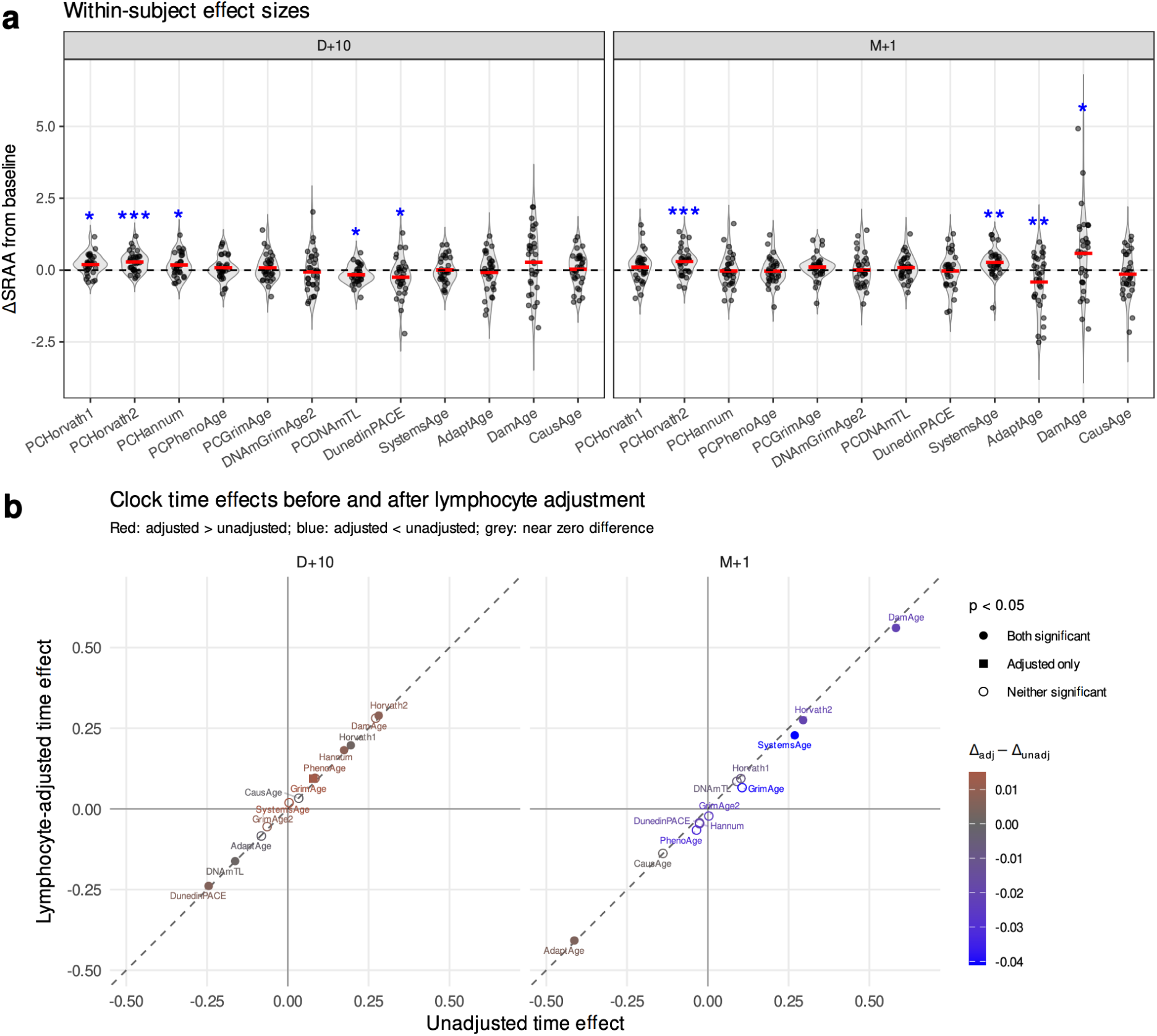
Sensitivity of clock time effects to lymphocyte adjustment. **a)** Within-subject changes from baseline in standardized residual age acceleration (ΔSRAA) at D+10 and M+1. This analysis corresponds to Figure 4a but was performed **without lymphocyte adjustment**. Each dot represents one participant; red bars indicate mean change; dashed lines mark no change from baseline. Blue asterisks indicate significant contrasts versus baseline from linear mixed-effects models including sex as a fixed-effect and subject as a random effect (*P < 0.05, **P < 0.01, ***P < 0.001). **b)** Comparison of component time-effect estimates before and after lymphocyte adjustment. The x-axis shows unadjusted estimates and the y-axis shows lymphocyte-adjusted estimates. The dashed diagonal indicates identical estimates. Color indicates the adjusted–unadjusted difference, and shape indicates significance status at P < 0.05.

**Supplementary Figure 8:**
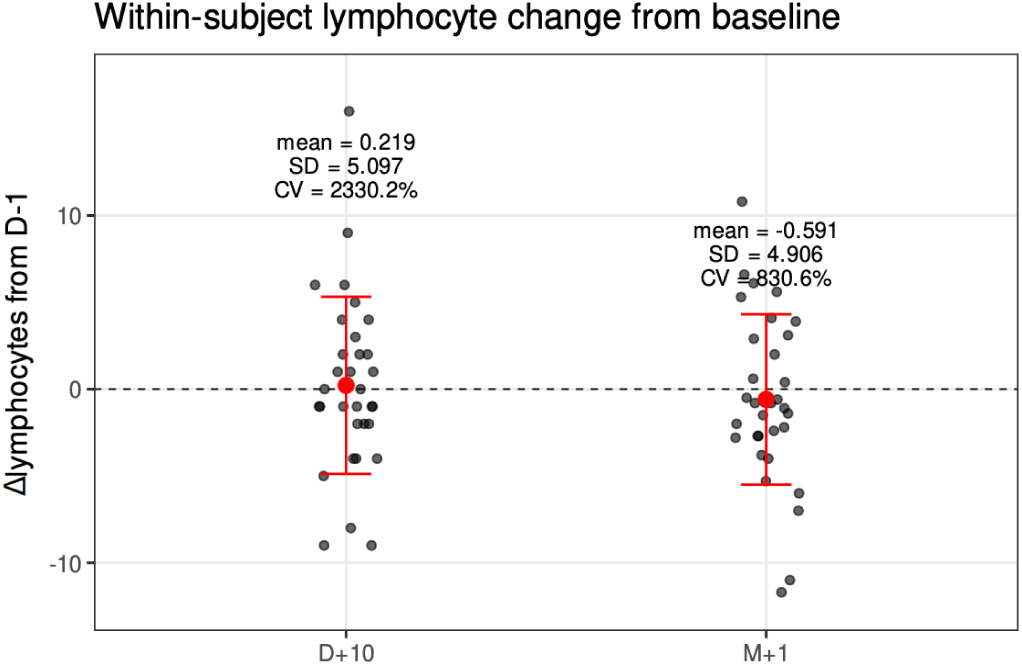
Within-subject lymphocyte changes from baseline at D+10 and M+1. Each black dot represents one participant. Red points and error bars indicate the mean change and ±1 standard deviation. The dashed horizontal line marks. Text annotations show the mean, standard deviation (SD), and coefficient of variation (CV).

**Supplementary Figure 9:**
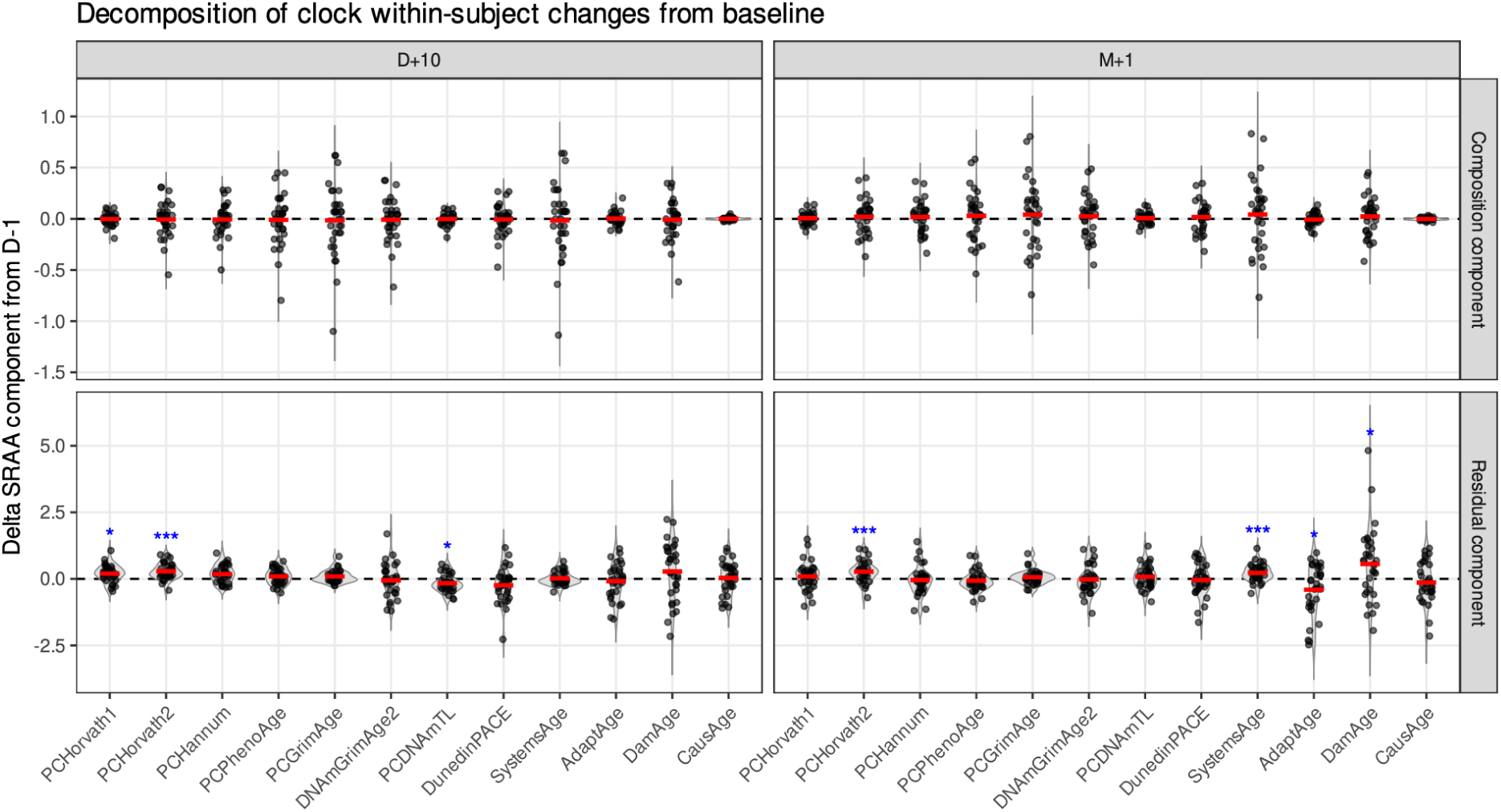
Decomposition of clock within-subject changes from baseline. For each epigenetic clock, within-subject ΔSRAA from baseline to D+10 and M+1 was decomposed into a lymphocyte composition component, shown in the top row, and a residual component, shown in the bottom row; see Methods. Plotting conventions are as in Supplementary Figure 7a.

**Supplementary Figure 10:**
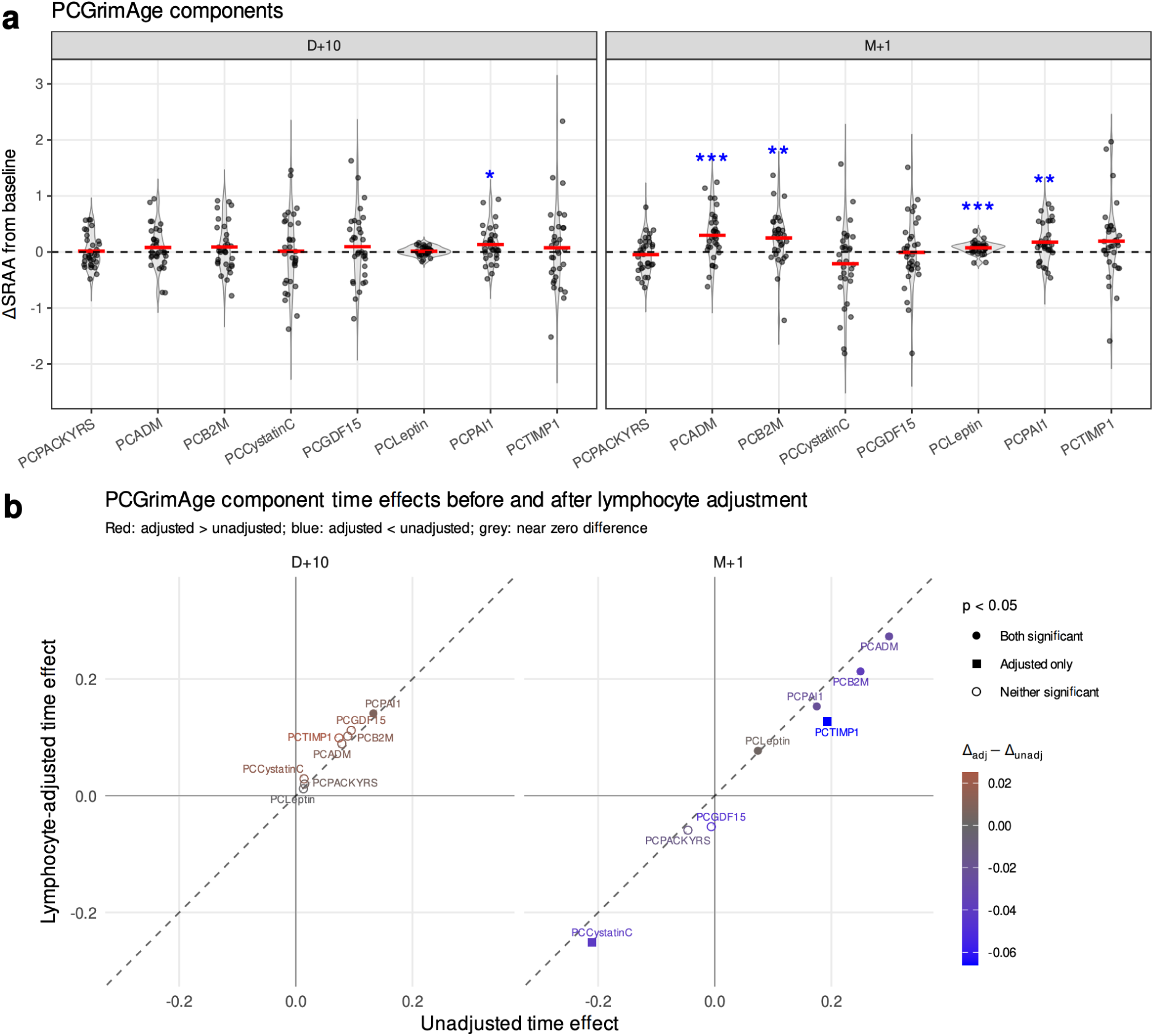
Sensitivity of PCGrimAge component time effects to lymphocyte adjustment. **a)** Within-subject changes from baseline in PCGrimAge component SRAA (ΔSRAA) at D+10 and M+1. This analysis corresponds to Figure 4b but was performed **without lymphocyte adjustment**. Each dot represents one participant; red bars indicate mean change; dashed lines mark no change from baseline. Blue asterisks indicate significant contrasts versus baseline from linear mixed-effects models including sex as a fixed effect and subject as a random effect (*P < 0.05, **P < 0.01, ***P < 0.001). **b)** Comparison of component time-effect estimates before and after lymphocyte adjustment. The x-axis shows unadjusted estimates and the y-axis shows lymphocyte-adjusted estimates. The dashed diagonal indicates identical estimates. Color indicates the adjusted–unadjusted difference, and shape indicates significance status at P < 0.05.

**Supplementary Figure 11:**
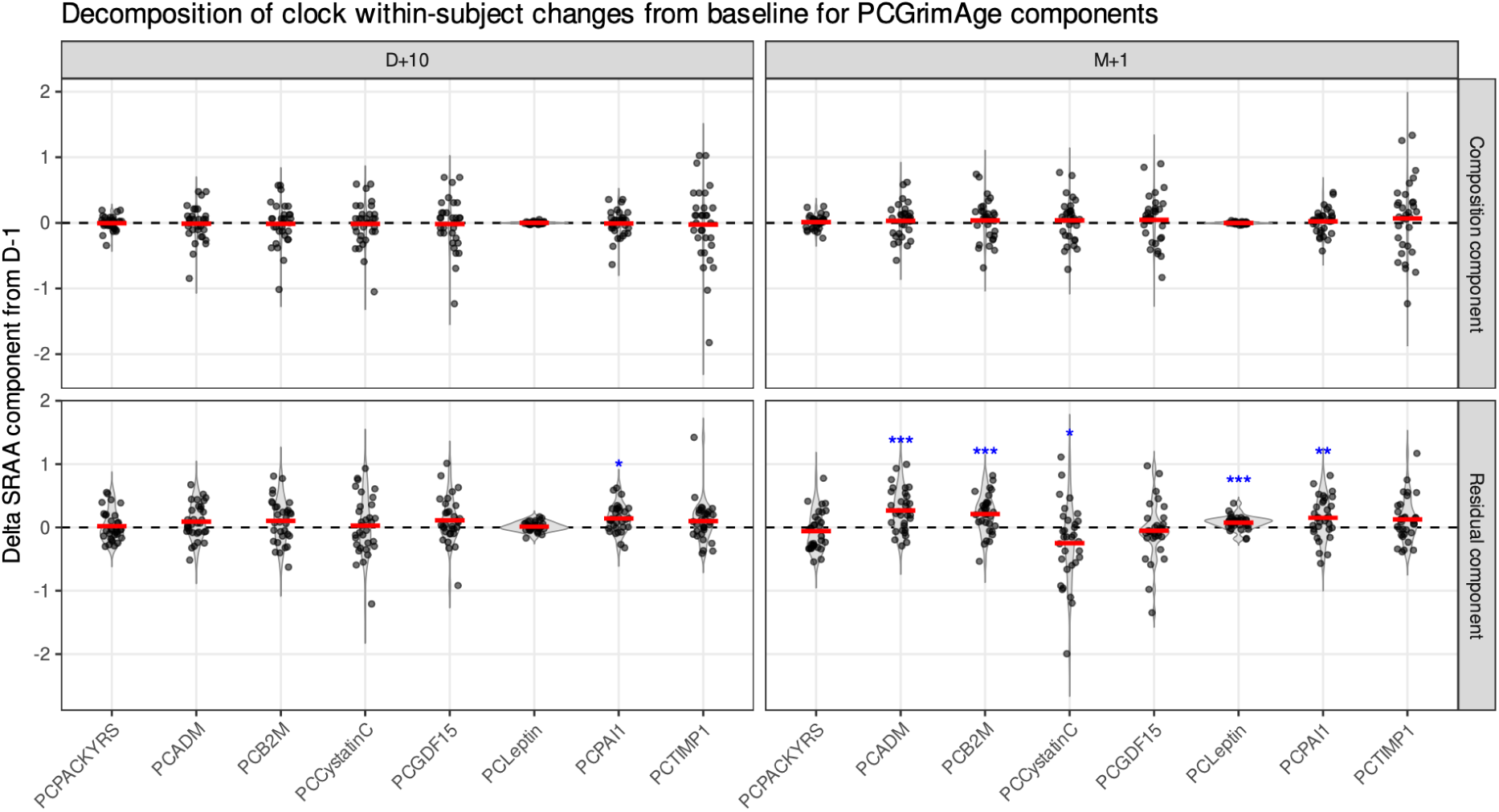
Decomposition of PCGrimAge component within-subject changes from baseline. For each PCGrimAge component, within-subject ΔSRAA from baseline to D+10 and M+1 was decomposed into a lymphocyte composition component, shown in the top row, and a residual component, shown in the bottom row; see Methods. Plotting conventions are as in Supplementary Figure 10a.

**Supplementary Figure 12:**
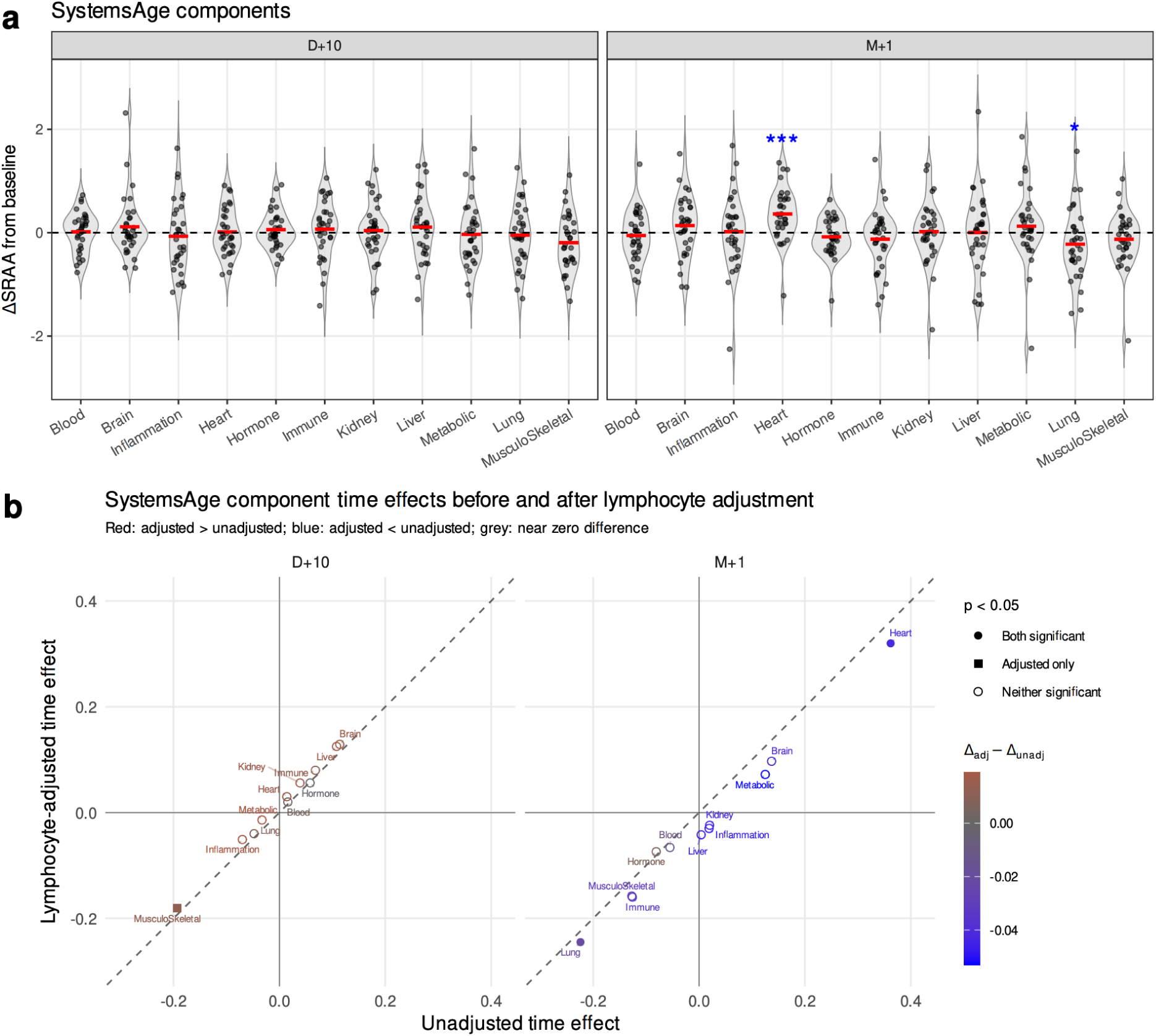
Sensitivity of SystemsAge component time effects to lymphocyte adjustment. **a)** Within-subject changes from baseline in SystemsAge component SRAA (ΔSRAA) at D+10 and M+1. This analysis corresponds to Figure 4c but was performed **without lymphocyte adjustment**. Each dot represents one participant; red bars indicate mean change; dashed lines mark no change from baseline. Blue asterisks indicate significant contrasts versus baseline from linear mixed-effects models including sex as a fixed effect and subject as a random effect (*P < 0.05, **P < 0.01, ***P < 0.001). **b)** Comparison of component time-effect estimates before and after lymphocyte adjustment. The x-axis shows unadjusted estimates and the y-axis shows lymphocyte-adjusted estimates. The dashed diagonal indicates identical estimates. Color indicates the adjusted–unadjusted difference, and shape indicates significance status at P < 0.05.

**Supplementary Figure 13:**
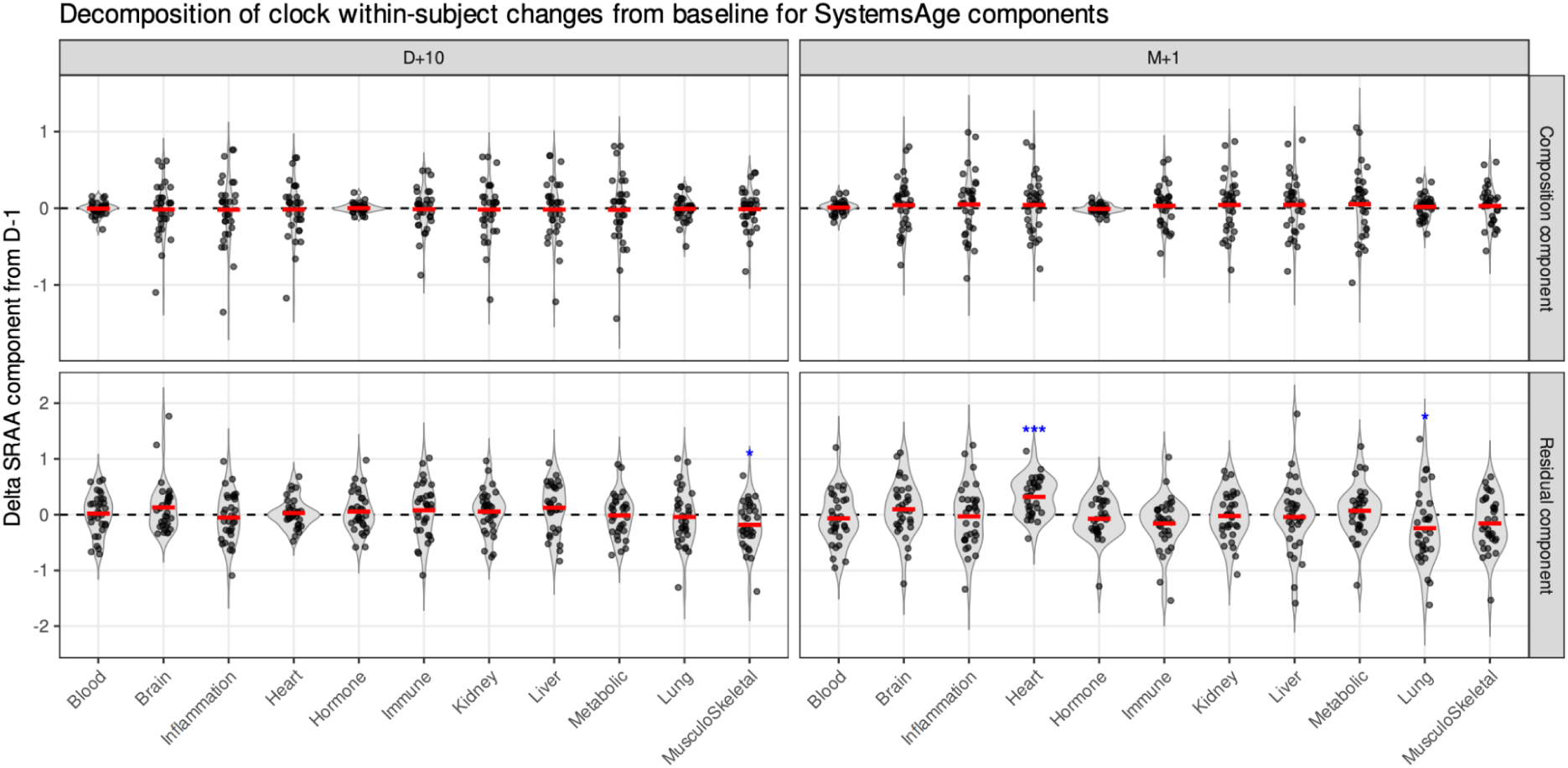
Decomposition of SystemsAge component within-subject changes from baseline. For each SystemsAge component, within-subject ΔSRAA from baseline to D+10 and M+1 was decomposed into a lymphocyte composition component, shown in the top row, and a residual component, shown in the bottom row; see Methods. Plotting conventions are as in Supplementary Figure 12a.

**Supplementary Figure 14:**
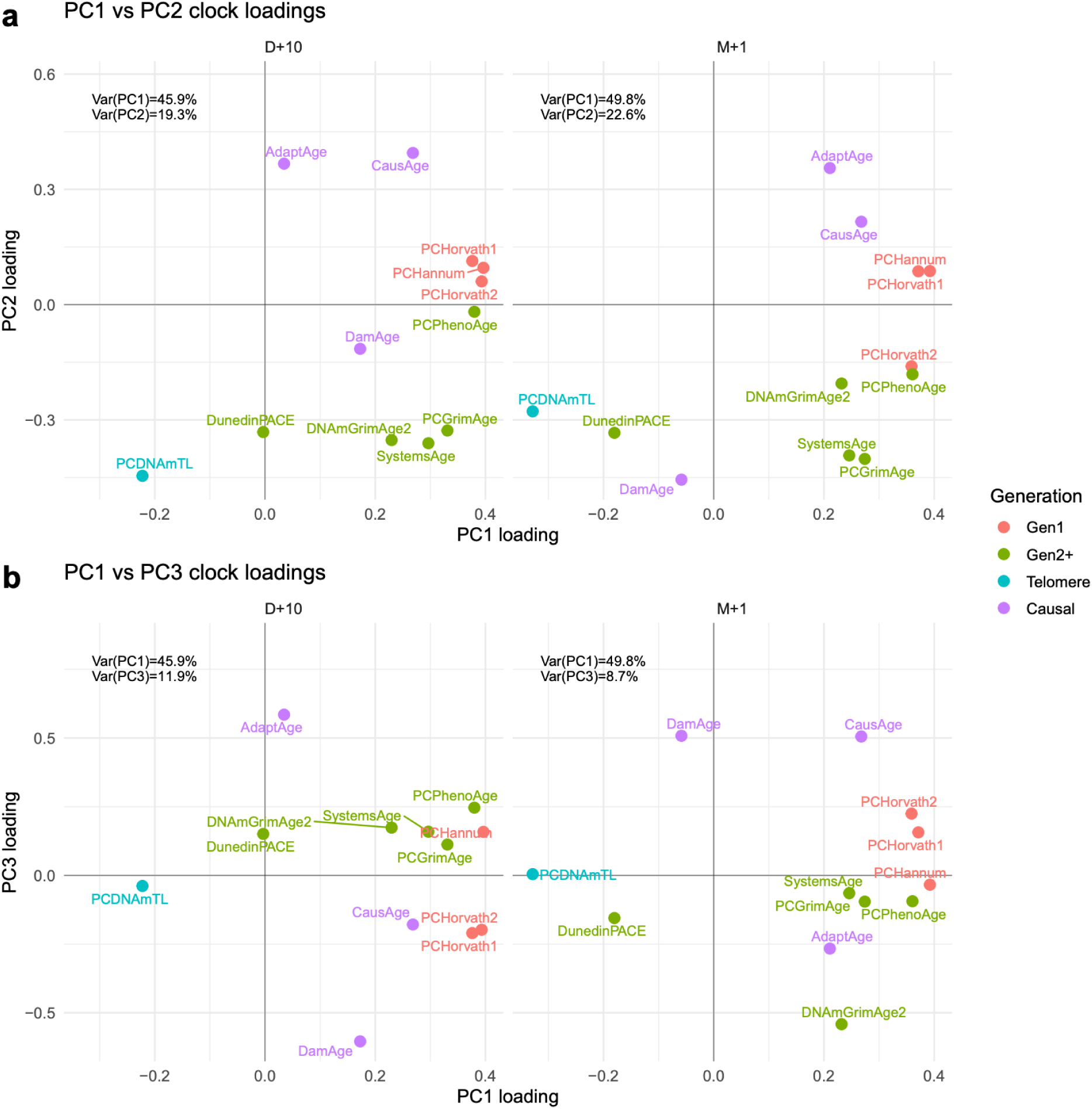
Principal component analysis (PCA) of within-subject ΔSRAA values at D+10 and M+1. Clock loading structures are shown separately for each time point. Panel **a)** displays the PC1–PC2 loading space, and panel **b)** displays the PC1–PC3 loading space. Each point represents an epigenetic clock, with colors indicating clock generation or conceptual class.Percentages indicate the proportion of variance explained by each principal component.

**Supplementary Figure 15:**
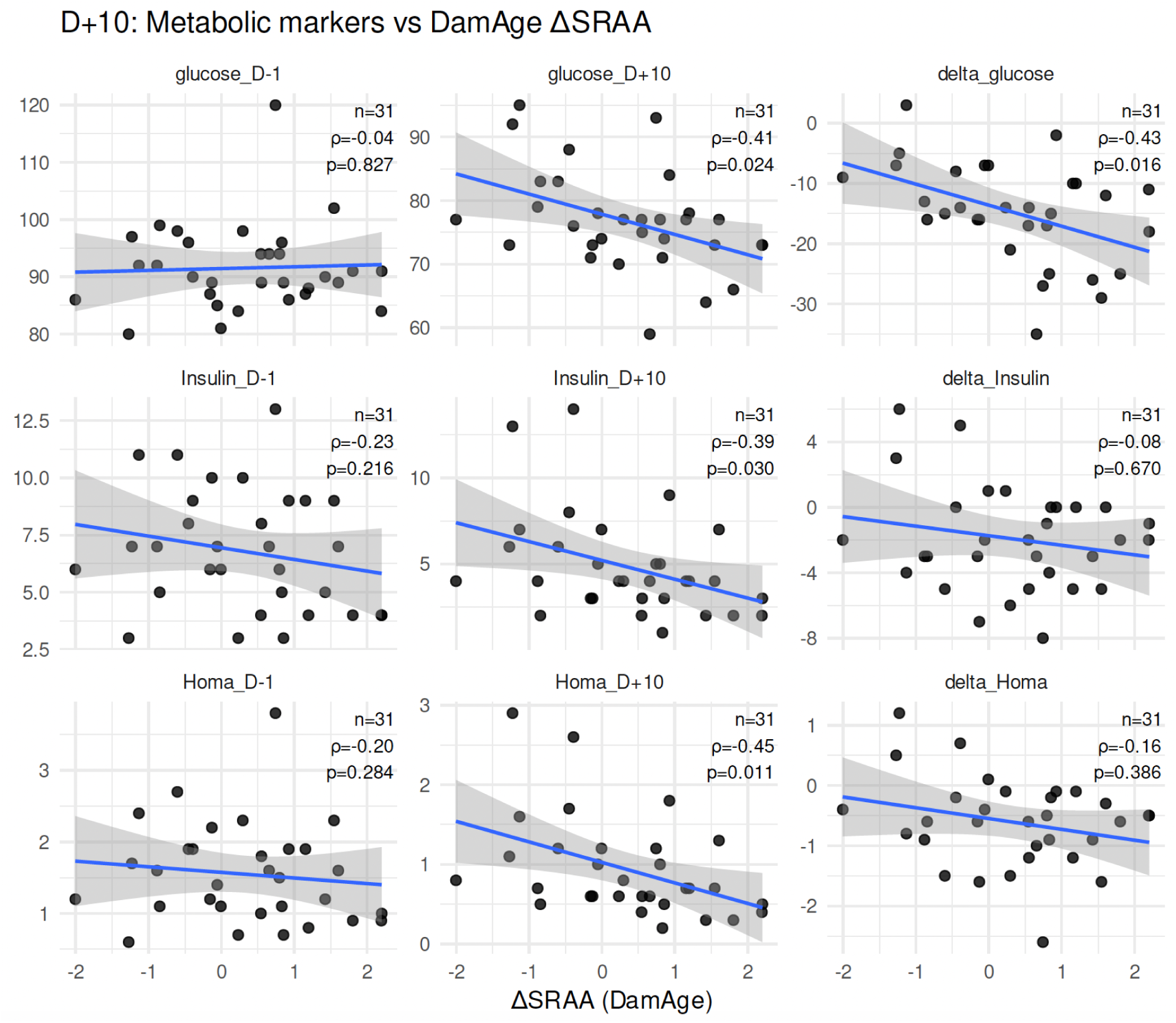
Associations between metabolic markers and DamAge responses during fasting. Scatter plots showing associations between standardized residual age acceleration derived from DamAge (ΔSRAA) and metabolic markers (glucose, insulin, and HOMA index) at baseline (D−1), during fasting (D+10), and as change-from-baseline (Δ) measures. Each point represents an individual participant (n = 31). Blue lines indicate fitted linear trends with shaded areas representing 95% confidence intervals. Spearman correlation coefficients (ρ) and corresponding p-values are shown in each panel.

**Supplementary Figure 16:**
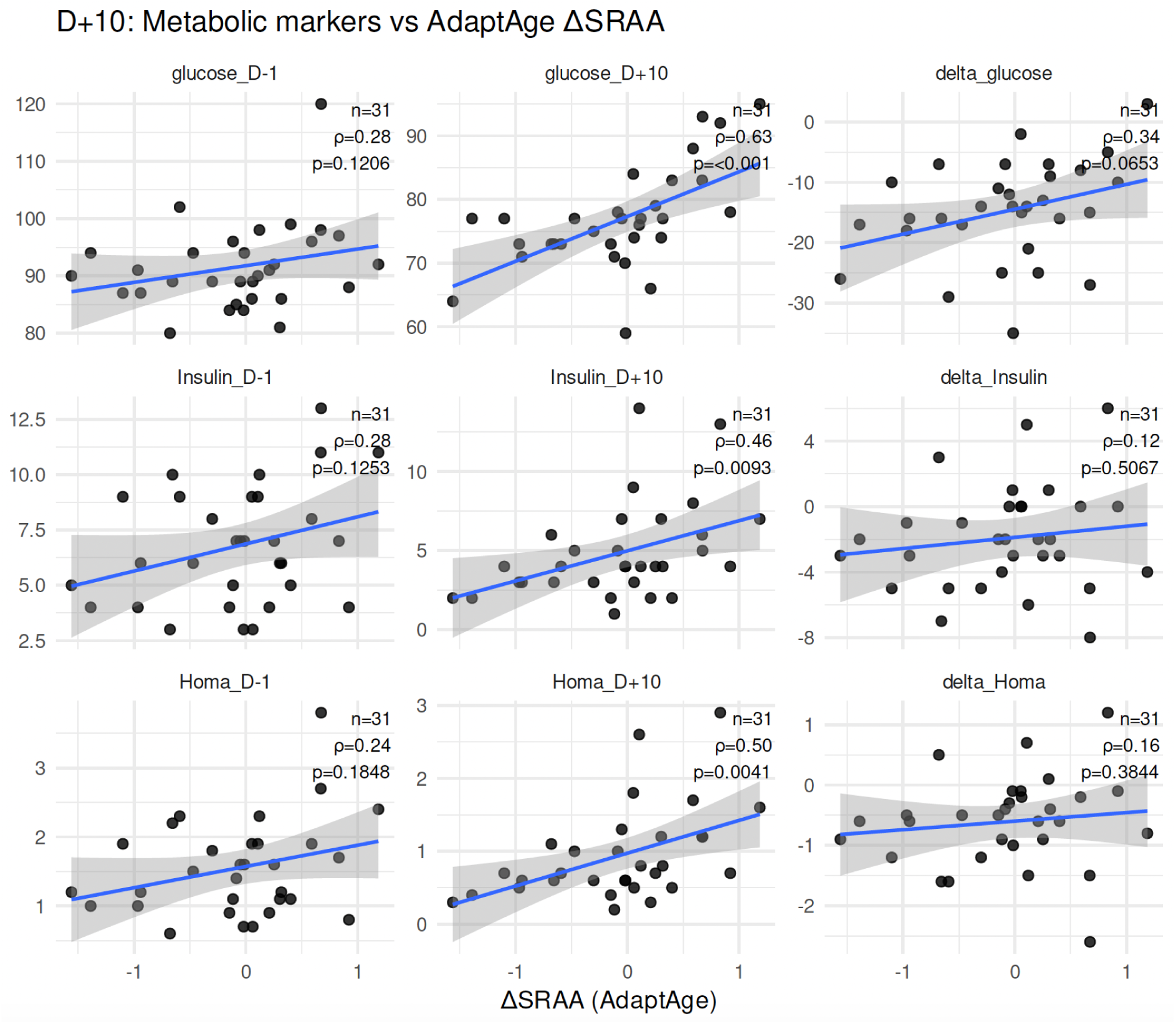
Associations between metabolic markers and AdaptAge responses during fasting. Same analysis as Supplementary Figure 15, showing associations between AdaptAge standardized residual age acceleration (ΔSRAA) and metabolic markers (glucose, insulin, and HOMA) at baseline (D−1), during fasting (D+10), and as change-from-baseline (Δ) values. Each point represents an individual participant (n = 31). Blue lines indicate linear fits with shaded 95% confidence intervals. Spearman correlation coefficients (ρ) and p-values are shown in each panel.

**Supplementary Figure 17:**
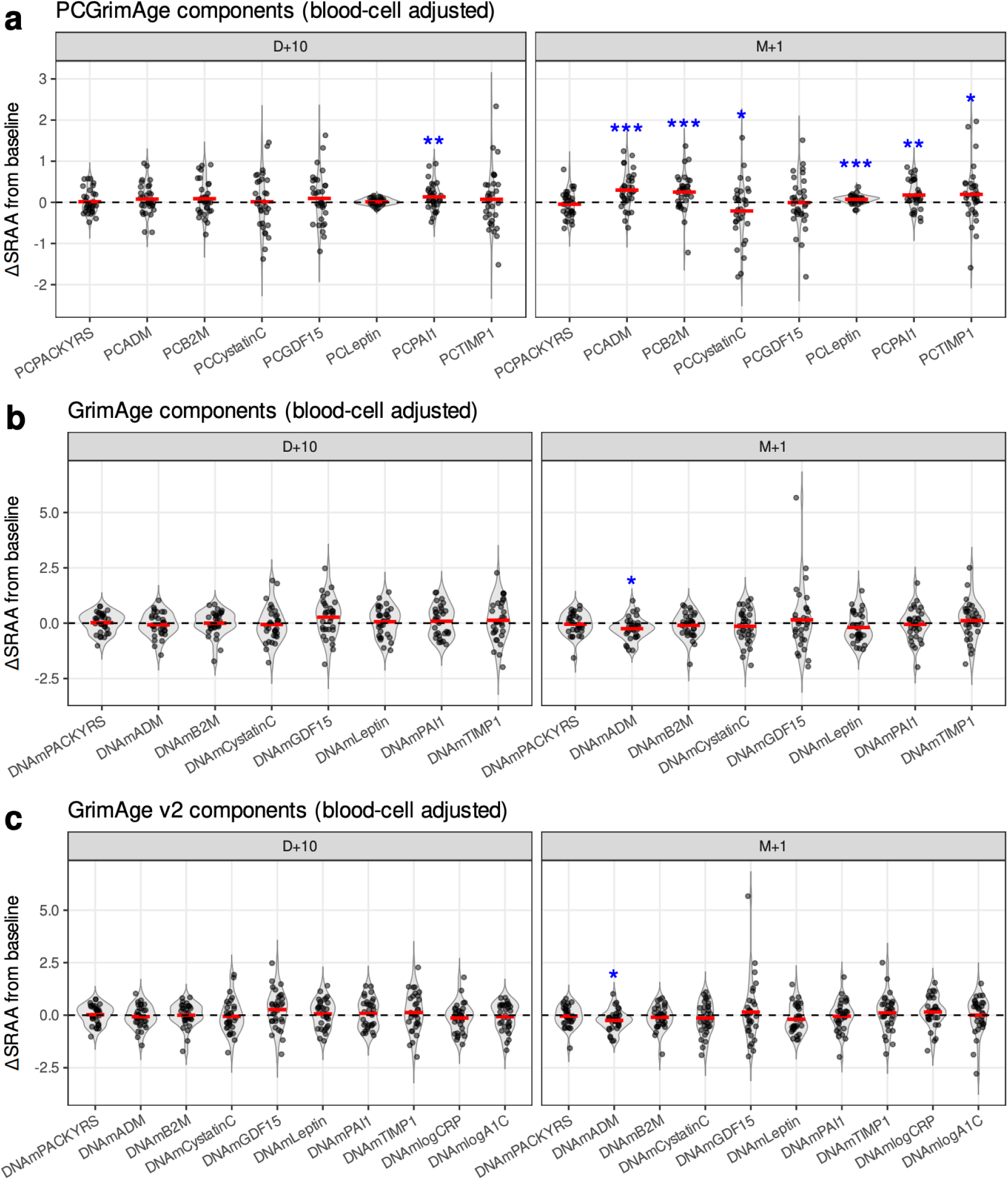
Comparison of ΔSRAA for PCGrimAge, GrimAge, and GrimAge v2. Individual DNAm surrogate components are shown for each clock implementation. Red bars indicate group means; dashed lines indicate no change from baseline. Asterisks denote statistically significant deviations from baseline (same test as in Figure 4).

**Supplementary Figure 18:**
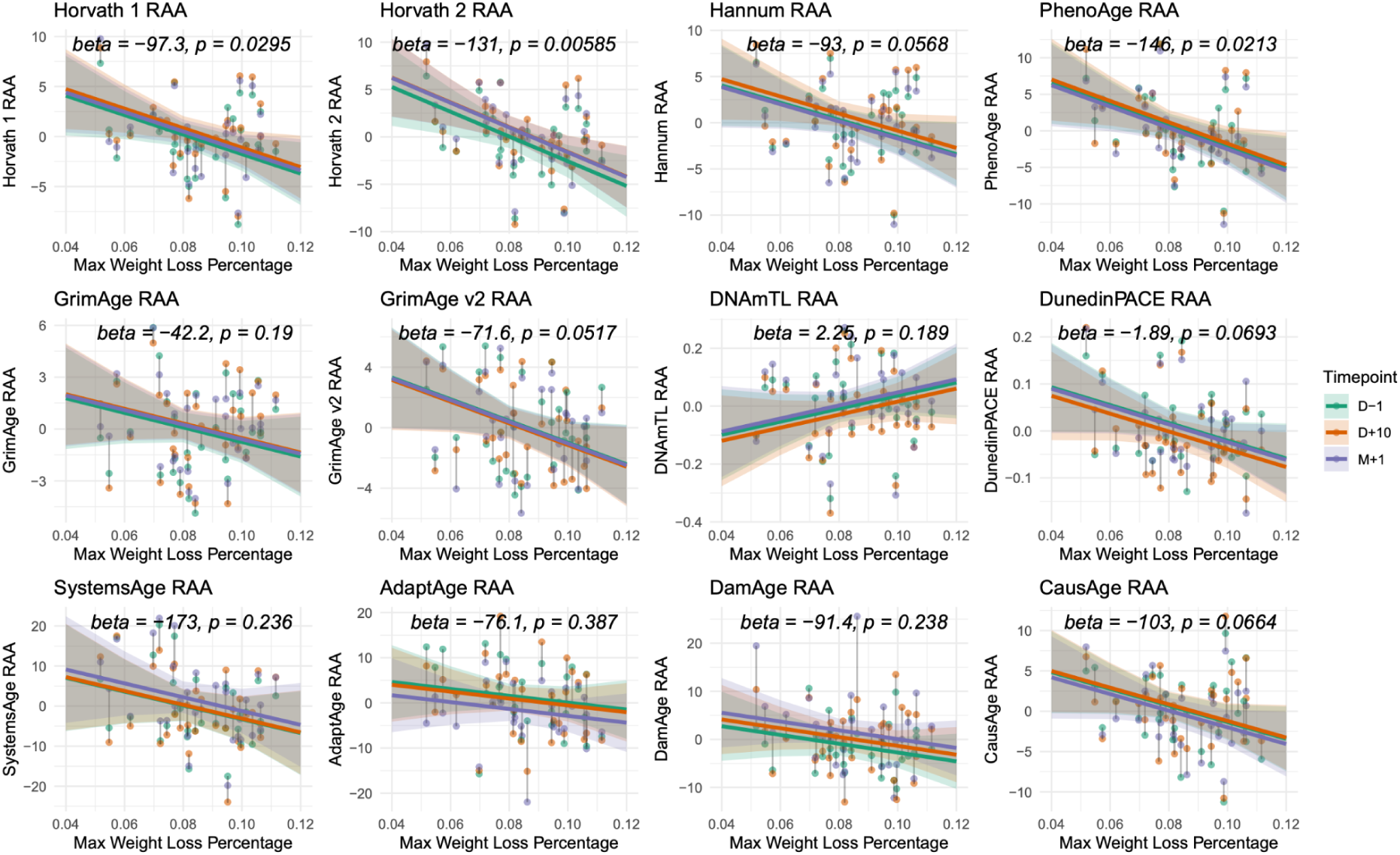
Association between residual epigenetic age acceleration and maximum weight loss percentage across 12 clocks (without BMI adjustment). This figure extends the analysis shown for six representative clocks in Figure 5 to all 12 clocks, using the same model specification and without baseline BMI adjustment. Residual age acceleration was modeled using linear mixed-effects models including maximum weight loss percentage, time point, sex, and lymphocyte proportions as fixed effects, with participant ID as a random effect. Plotting conventions are as in Figure 5: the marginal effect of maximum weight loss percentage at each time point is shown as a colored regression line, with the corresponding regression coefficient (beta) and p-value (p) displayed in each panel. Data points are colored by time points and connected by lines to indicate repeated measures from the same individual.

**Supplementary Figure 19:**
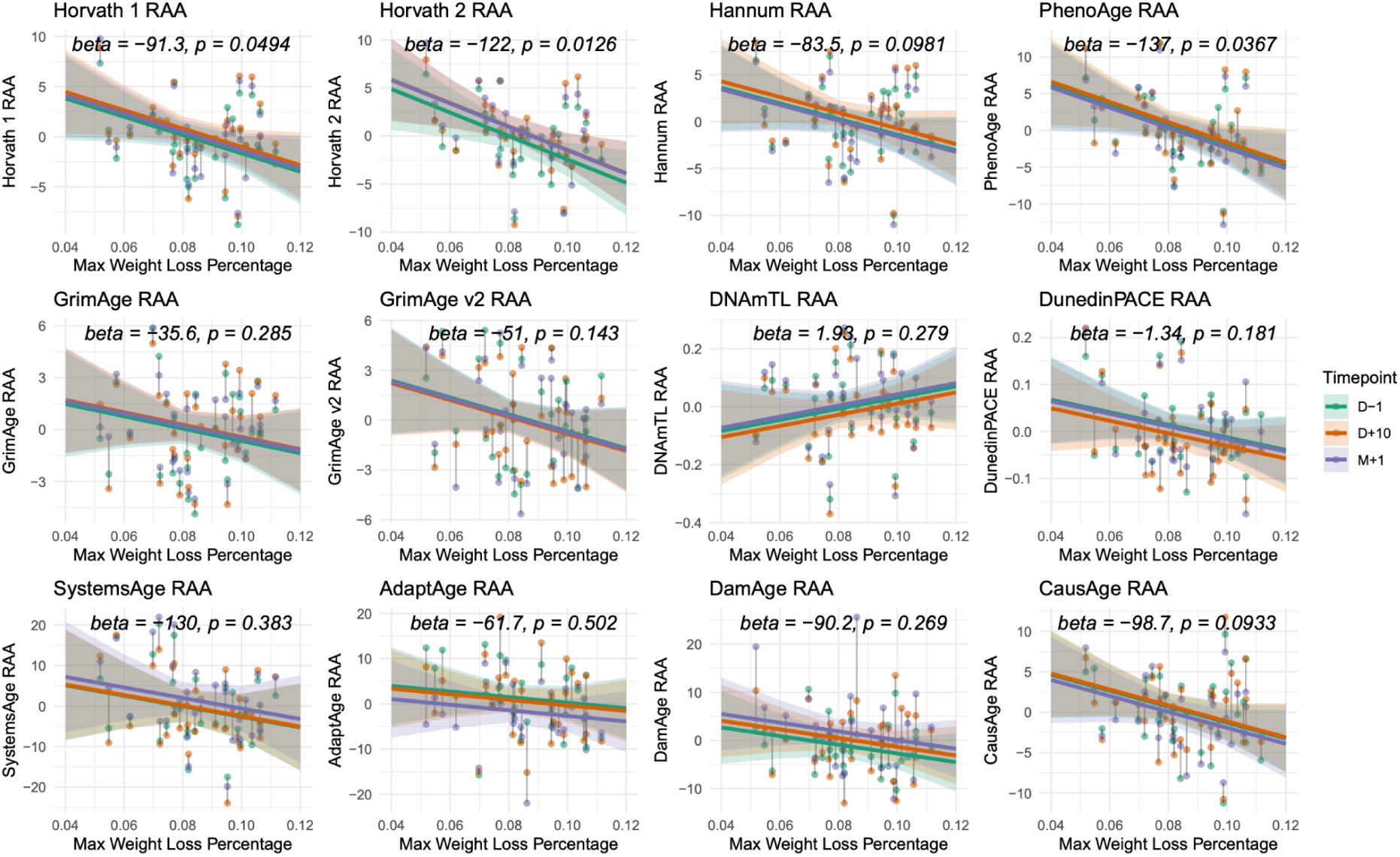
Association between residual epigenetic age acceleration and maximum weight loss percentage across 12 clocks (with baseline BMI adjustment). This figure shows the BMI-adjusted sensitivity analysis corresponding to Figure 5 and Supplementary Figure 18. Residual age acceleration was modeled using linear mixed-effects models including maximum weight loss percentage, time point, sex, lymphocyte proportions and **baseline BMI** as fixed effects, with participant ID as a random effect. Plotting conventions are as in Figure 5 and Supplementary Figure 18.

## Supplementary tables

**Supplementary Table 1:** Vital parameters measured at D-1, D+4, D+10 and M+1. Values are presented as mean ± SD. Significance levels for comparisons with D–1 are indicated as follows: * = p<0.05, ** = p<0.01, *** = p<0.001. **** = p<0.0001. Not all parameters were measured at M+1 (indicated with NA).

| Parameter | D-1 | D+4 | D+10 | M+1 |
| --- | --- | --- | --- | --- |
| Weight (kg) | 79.16 ± 13.01 | 76.19 ± 12.50 **** | 72.94 ± 12.00 **** | 75.67 ± 12.12 **** |
| Body mass index (kg/m<sup>2</sup>) | 26.14 ± 3.92 | 25.16 ± 3.84 **** | 24.12 ± 3.89 **** | 25.21 ± 3.64 **** |
| Resting heart rate (bpm) | 67.09 ± 13.80 | 68.97 ± 12.22 | 67.88 ± 11.77 | NA |
| Systolic blood pressure (sbp) (mmHg) | 124.19 ± 15.40 | 121.82 ± 14.86 | 114.44 ± 13.53 ** | NA |
| Diastolic blood pressure (dbp) (mmHg) | 79.65 ± 10.69 | 74.28 ± 13.05 *** | 72.33 ± 8.55 ** | NA |

## Notes

### Competing Interest Statement

Authors M.K.. F.G., F.W.T. and R.M. are employees of the Buchinger Wilhelmi Development and Holding GmbH, Ueberlingen. CK, LC, SN, SGN are employees of GenKnowme. The Regents of the University of California are the sole owner of patents and patent applications directed at epigenetic biomarkers for which Steve Horvath is a named inventor. S.H. is a founder and paid consultant of the non-profit Epigenetic Clock Development Foundation that licenses these patents. The remaining authors declare that the research was conducted in the absence of any commercial or financial relationships that could be construed as a potential conflict of interest.

